# Brain signal predictions from multi-scale networks using a linearized framework

**DOI:** 10.1101/2022.02.28.482256

**Authors:** Espen Hagen, Steinn H. Magnusson, Torbjørn V. Ness, Geir Halnes, Pooja N. Babu, Charl Linssen, Abigail Morrison, Gaute T. Einevoll

## Abstract

Simulations of neural activity at different levels of detail are ubiquitous in modern neurosciences, aiding the interpretation of experimental data and underlying neural mechanisms at the level of cells and circuits. Extracellular measurements of brain signals reflecting transmembrane currents throughout the neural tissue remain commonplace. The lower frequencies (≲ 300Hz) of measured signals generally stem from synaptic activity driven by recurrent interactions among neural populations and computational models should also incorporate accurate predictions of such signals. Due to limited computational resources, large-scale neuronal network models (≳ 10^6^ neurons or so) often require reducing the level of biophysical detail and account mainly for times of action potentials (‘spikes’) or spike rates. Corresponding extracellular signal predictions have thus poorly accounted for their biophysical origin.

Here we propose a computational framework for predicting spatiotemporal filter kernels for such extracellular signals stemming from synaptic activity, accounting for the biophysics of neurons, populations, and recurrent connections. Signals are obtained by convolving population spike rates by appropriate kernels for each connection pathway and summing the contributions. Our main results are that kernels derived via linearized synapse and membrane dynamics, distributions of cells, conduction delay, and volume conductor model allow for accurately capturing the spatiotemporal dynamics of ground truth extracellular signals from conductance-based multicompartment neuron networks. One particular observation is that changes in the effective membrane time constants caused by persistent synapse activation must be accounted for.

The work also constitutes a major advance in computational efficacy of accurate, biophysics-based signal predictions from large-scale spike and rate-based neuron network models drastically reducing signal prediction times compared to biophysically detailed network models. This work also provides insight into how experimentally recorded low-frequency extracellular signals of neuronal activity may be approximately linearly dependent on spiking activity. A new software tool `LFPykernels` serves as a reference implementation of the framework.

**Author summary:** Understanding the brain’s function and activity in healthy and pathological states across spatial scales and times spanning entire lives is one of humanity’s great undertakings. In experimental and clinical work probing the brain’s activity, a variety of electric and magnetic measurement techniques are routinely applied. However interpreting the extracellularly measured signals remains arduous due to multiple factors, mainly the large number of neurons contributing to the signals and complex interactions occurring in recurrently connected neuronal circuits. To understand how neurons give rise to such signals, mechanistic modeling combined with forward models derived using volume conductor theory has proven to be successful, but this approach currently does not scale to the systems level (encompassing millions of neurons or more) where simplified or abstract neuron representations typically are used. Motivated by experimental findings implying approximately linear relationships between times of neuronal action potentials and extracellular population signals, we provide a biophysics-based method for computing causal filters relating spikes and extracellular signals that can be applied with spike times or rates of large-scale neuronal network models for predictions of population signals without relying on *ad hoc* approximations.

## Introduction

Extracellular electric recordings of neuronal activity, either by embedding sharp electrodes in neural tissue [1] or by placing electrodes on top of cortex [2] or on the scalp (electroencephalography – EEG [3]), have a long history in the experimental and clinical neurosciences. The same applies to magnetic recordings outside of the head (magnetoencephalography – MEG [4]). However, the link between the measured brain signals and the underlying neuronal activity remains poorly understood due to the inherent ill-posed inverse problem: The number of contributing sources is large compared to the limited number of discrete locations in- and outside of the brain tissue where one can measure. However, the forward problem is well-posed, given the transmembrane currents in all neurons setting up the activity. Different electric and magnetic signals can be computed by means of so-called volume conductor (VC) theory mapping source currents to each signal type, thus models accounting for the biophysical properties of neurons and networks thereof can now be used to study the link between activity and measurements [5, 6].

Dynamics of biophysically detailed neurons and synaptically coupled networks thereof are typically modeled by solving sets of coupled, linear and non-linear ordinary or partial differential equations describing the dynamics of the neuronal membranes, ion channel conductances, synapses, and so forth (see e.g., [7]). Multicompartment (MC) models have for decades been the go-to tool for geometrically detailed conductance-based neuron models as tailored software solvers are readily available such as NEURON [8], GENESIS [9], and Arbor [10]. For the purpose of computing extracellular electric and magnetic signals, transmembrane currents from the MC neuron simulation are then combined with the appropriate forward model derived using linear volume conductor theory, as incorporated in software interfacing the neural simulator like LFPy [11, 12], NetPyNe [13], and BMTK [14]. For brain tissues, a linear relationship between transmembrane currents and extracellular electric potentials as well as magnetic fields appears well established [3, 15–17].

Illustrated in Fig 1A, neuronal network models may account for different levels of detail, ranging from biophysically detailed MC neuron networks (top level), simplified spiking point-neuron networks (middle level) and population type models accounting for population-averaged activity (bottom level). The different levels may at times be bridged with appropriate mapping of parameters. As illustrated, MC models may be directly combined with VC theory for extracellular signal predictions as these models account for the spatiotemporal distribution of transmembrane currents, while the less detailed models, in particular point-neuron networks and mean-field type population models, do not. Thus in order to relate their activity in terms of spike times or spike rates of the different populations to extracellular signals additional steps are required, here illustrated by some ‘black box’ model taking spikes or equivalent spike rates of each population as input while outputting approximated extracellular signals.

**Fig 1.**
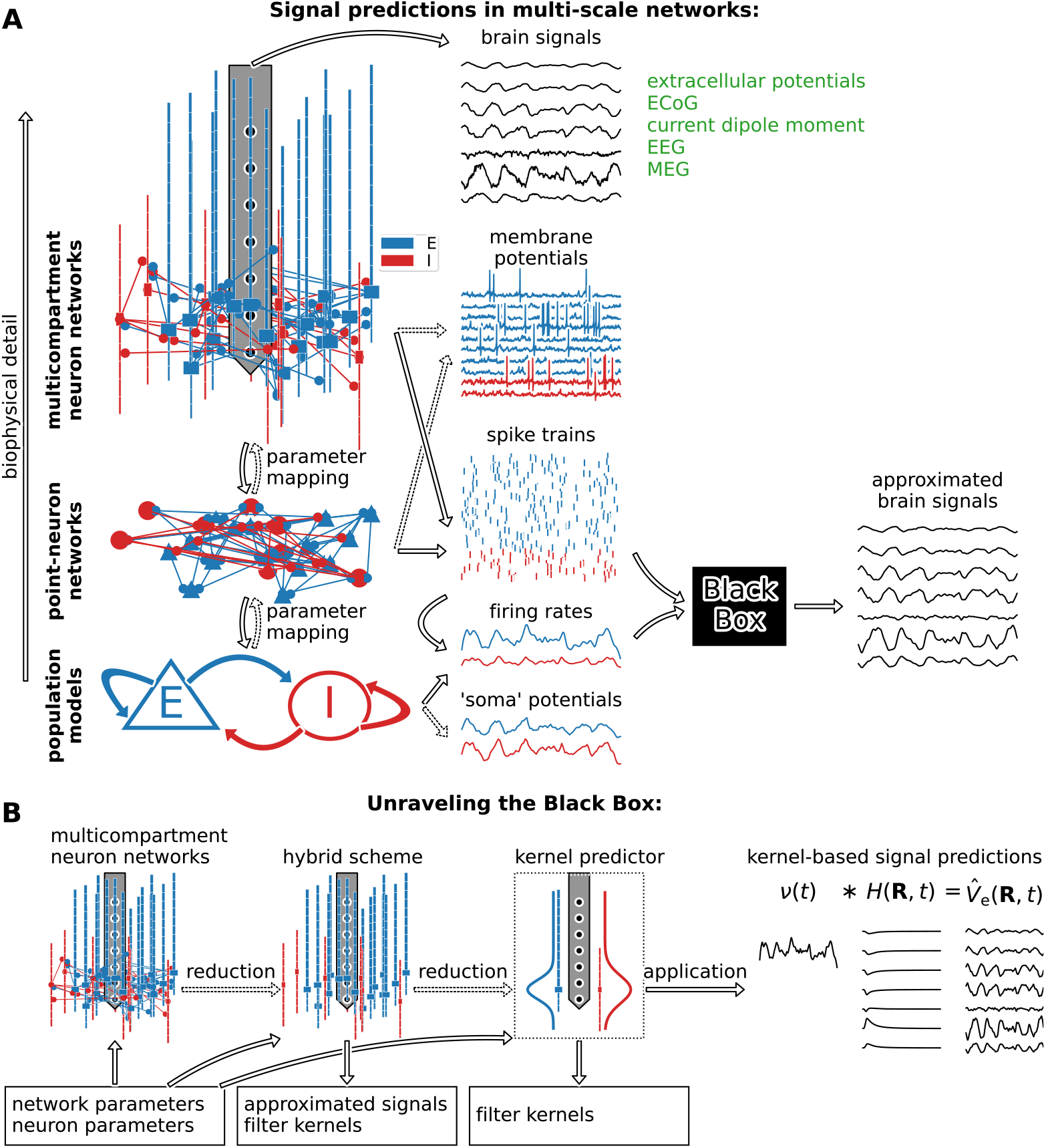
Levels of detail for neuronal network models and roadmap for approximated brain signal predictions. (**A**) Biophysically detailed MC neuron network models at the microscopic scale allow for simulating synaptic connectivity and whole-cell dynamics, including APs, spike trains and extracellular signals (e.g., the extracellular potential) using forward models derived via VC theory. Neither less detailed spiking point-neuron network models nor continuous population type models (neural mass models, mean-field models, neural field models) towards the mesoscopic scale facilitate extracellular signal predictions. They require a model translating spike events or spike rates into representative extracellular signal approximation, here illustrated by the black box. (**B**) (1) Detailed networks provide ground truth signals and spiking activity for successive reduction steps, given a set of neuron and network parameters (box). (2) The ‘hybrid scheme’ setup [18] relies on simulating MC neuron populations but omits recurrent network connections. Predictions are governed by spike times of recurrent networks and may use linearized neural dynamics. (3) The ‘kernel predictor’ setup relies on a subset of MC neuron simulations and accounts for the underlying statistics (synapse densities, etc.) of the network, and computes spatiotemporal spike-to-signal impulse response functions. (4) Firing rates of presynaptic populations *v*(*t*) are convolved by precomputed kernels *H*(**R**, *τ*) for signal approximations.

Illustrated in Fig 1B, we shall approach this black box problem by models that account for key properties of the biophysically detailed network models (mainly cell model membrane dynamics, spatial distributions of cells and synapses, network connectivity, temporal synapse dynamics), properties which could also be constrained by experimental data. Through a systematic reductionistic approach we first apply the so-called ‘hybrid scheme’ for extracellular signal predictions [18] which entails that presynaptic spike events are first simulated in the actual network in Fig 1B. The spike times are used for synapse activation times in corresponding populations of MC neurons as they would occur in the actual network. We shall approximate synaptic and ion-channel conductances by linearized variants as this allows for simulating approximated extracellular signals using fully linear models. The hybrid scheme predictions are validated against ground truth signals predicted by the true network models. The setup is also used to compute averaged causal spike-signal impulse response functions, ‘kernels’, for each connection pathway. Such hybrid scheme kernels shall be applied with population firing rates to approximate extracellular signals [18, 19]. Expanding on this kernel-based scheme, we here present a novel method to efficiently compute such kernels directly accounting for the biophysics and description of the putative network model and neurons. Their prediction relies on the same linearization steps introduced for the hybrid scheme but may bypass the hybrid scheme altogether. All kernel-based signal predictions are validated against corresponding ground truth signals. The computational schemes investigated here are applicable for predictions of the low-frequency content (≲ 300 Hz or so) of the signals usually associated with population activity and network interactions. The final kernel-based predictions can also readily be combined with spike- and rate-based network simulation frameworks.

In the above context, we in part consider observations and assumptions of near-linear relationships between times of neuronal action potentials (APs) as well as their extracellularly recorded correlates (‘spikes’), and low-frequency parts (below a few hundred hertz) of extracellularly recorded population signals like local field potential (LFP), EEG, and MEG signals [20–23]. For synaptically coupled neuronal networks, one may consider two main direct neuronal contributors to extracellular population signals. The first is due to presynaptic neurons generating APs observed as extracellular spikes nearby each active neuron in recordings using invasive microelectrodes. The second is due to evoked synaptic currents and associated membrane currents throughout postsynaptic populations following presynaptic APs. AP durations are on the order of milliseconds, APs occur with relative sparsity (low spike rates) and irregularity in single-neuron spike trains [24], observed pairwise spike train correlations are on average weak [25, 26], and extracellular spike amplitudes decay quickly with distance [27, 28]. Extracellular spikes also carry more power toward higher frequencies [27]. The latter synaptic contributions can be small in amplitude per pair of pre- and postsynaptic neurons relative to currents related to each presynaptic AP itself, but each neuron typically targets many neurons via hundred or even thousands of synapses, and recurrent interactions may affect the times of subsequent activations across large neural populations. The dynamics of synapse currents are also relatively slow and can thus be assumed to shape extracellular signals around lower frequencies than presynaptic contributions. We may then assume that mainly synaptic activity governs the low-frequency content of extracellular signals, in part, via a boosting effect on the compound signals by even weak pairwise spike train correlations [18].

Dynamics of neuronal activity are typically nonlinear, one prime example is the model for APs by [29] which also provided a mathematical formalism that remains commonly used to describe different ion-channel dynamics (e.g., [30]). Extracellularly recorded postsynaptic responses following presynaptic AP events can not initially be assumed to be linear, as synaptic currents following activation are not linearly dependent on the synaptic conductance due to membrane potential changes. Furthermore, there may be active (voltage- and calcium-dependent) ion channels present across dendrites resulting in non-linear integration even below AP threshold, and contributions by different activations of multiple synapses may not sum linearly [31]. Synaptic activity may also result in dendritic Ca^2+^- and NMDA spikes [32].

Thus to explain experimental data implying approximately linear relationships between times of presynaptic spikes and different electric signals, the direct signal contribution by both pre- and postsynaptic activity must sum approximately linearly. Furthermore, for synaptic currents across postsynaptic populations the different contributions by nonlinear synapse and membrane dynamics must be negligible or well explained by linear components around typical working points (e.g., average membrane potentials and spike rates). Still, a number of computational studies assume linearity between presynaptic spike events and corresponding times of synaptic activations and resulting extracellularly recorded signals [18, 19, 33–38]. Others assume linearity between transmembrane input current and extracellular potentials [39], in part justified by model work wherein dynamics of active ion channels are approximated by linear dynamics [40, 41]. Analyses of experimental recordings by [42] also show synaptic currents and the LFP to be strongly coupled using a linear regression model.

Henceforth, we shall examine the validity of models that either explicitly or implicitly assume linear relationships between neuronal spiking and extracellular signals. We will do so by comparing the extracellular signals that these models predict with corresponding predictions obtained with biophysically detailed MC neuron networks. Hereby we test the following approaches (hypotheses): (1) Linearized model setups can accurately capture the spatiotemporal features of ground truth extracellular potential and current dipole moment computed from recurrent networks of inherently non-linear constituents. For this testing, we first apply the hybrid prediction scheme [18]. (2) If the linear hybrid scheme implementation accurately captures the ground truth signals, we test whether or not the output extracellular signal predictions can be well captured as a linear and time-invariant causal system, taking population spike rates as input filtered by suitable spatiotemporal causal filters. These sets of filters or ‘kernels’ represent postsynaptic spike-signal impulse responses averaged over pairs of pre- and postsynaptic populations, and are initially computed via the hybrid scheme. (3) Knowledge of the underlying distributions of cells and synapses, conduction delays, linearized cell, and synapse dynamics, and corresponding population spike rates is sufficient information to predict these spatiotemporal causal kernels accurately.

The kernel-based approach can be applied with recurrent neuronal network descriptions using much-simplified neuron representations, like leaky integrate-and-fire (LIF) point neurons, variants thereof, as well as few-compartment neuron models, as the main determinant for the extracellular signals is presynaptic spikes or spike rates. Also, point-neuron networks may accurately mimic experimentally observed spiking activity as well as corresponding MC neuron networks (see e.g., [43]). Then, the computationally costly MC simulations may only be required in order to compute the appropriate sets of kernels, thus reducing compute resource demands by orders of magnitude. This may for instance open for efficient forward-model-based extracellular signal predictions from large-scale point-neuron network models encompassing multiple brain areas [44] or models incorporating realistic cell densities within an area [36, 43]. The kernel methodology would also be immediately useful with rate-based frameworks, as also population spike rates of spiking network models may be accurately captured in corresponding population rate models (see e.g., [45–48]).

This study is organized as follows: In Materials and methods we first detail a generic biophysically detailed MC neuron network that is used for ground truth signal predictions, and different network configurations. Then we detail a proposed hybrid methodology that allows for separation between simulations of network activity (‘spikes’) and extracellular signals, and the derivation of linearized signal predictions, including our proposed methodology for fast, accurate, and deterministic predictions of kernels. In Results we investigate the properties of neuron models in active and linearized versions, recurrent MC neuron networks and compare the different linear approximations to the corresponding ground truth signals. Then, we showcase the kernel-based methodology to network spiking activity of a recurrent network of leaky integrate-and-fire neurons. In Discussion we consider the implications of this work and possible future steps.

## Materials and methods

### Reference multicompartment neuron networks

We first define the properties of a generic recurrently connected network of MC neurons used for ground truth signal generation and later signal approximations. For compactness, we choose a symbolic notation similar to [18] wherever possible and provide the model details as a generic ‘recipe’. Their particular values are summarized in this section and Tables 1 to 3. In general terms we:

1. Let *X* ∈ {…} and *Y* ⊆ *X* denote pre- and postsynaptic populations, respectively. Each population corresponds to separate classes of neurons (derived from anatomy, electrical properties, gene expression, phenomenology, etc.). We let populations in *Y* be a local subset to allow for remote neuronal populations in *X*. (Thus *X* may include remote populations, point processes, external stimuli, and similar, which we will assume give approximately zero direct contributions to the local signals predicted by the full recurrent network model).
2. Let lists *N*_*X*_ and *N*_*Y*_ denote the sizes of populations *X* and *Y*.
3. Let *u* ∈ {1, …, *N*|*N* ∈ *N*_*X*_} and *v* ∈ {1, …, *N*|*N* ∈ *N*_*Y*_} denote pre- and postsynaptic neuron indices, respectively.
4. Let 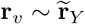 denote a discretely sampled somatic location of neuron *v*, where 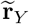 describes the probability density function of somatic locations of population *Y* in 3-dimensional (3D) space.
5. Let *K*_*Y X*_ denote the total number of pairwise connections between presynaptic (source) population *X* and postsynaptic (target) population *Y*. Assuming random connectivity with binomial in- and out-degree distributions the corresponding connection probability is then 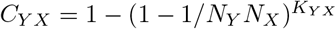 [49]. *K*_*Y X*_ ≈ *C*_*Y X*_*N*_*Y*_ *N*_*X*_ for small connection probabilities. The subscript *Y X* notation is used throughout this paper to emphasize that these parameters are connection-specific.
6. Let 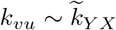 denote the randomly sampled number of synapses (multapses) per connection if a connection exist between neurons *u* and 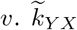 here describes a discrete distribution from which integer numbers greater than 0 are drawn.
7. Let probabilities of synapse placement onto postsynaptic compartments indexed by *m* be proportional to the product *L*_*Y X*_ (*z*_*m*_)*A*_*m*_, where *L*_*Y X*_ (*z*) is a depth-dependent function evaluated at the midpoint of each compartment with surface area *A*_*m*_. Compartments are indexed by *m*. Synapse placements are drawn randomly *k*_*vu*_ times for each pair of connected neurons.
8. Let the current for each synapse following activation at time *t* be described by

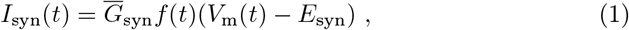

where 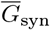 denotes the maximum synaptic conductance and *f* (*t*) ∈ [0, 1] the temporal kinetics of the synapse. We let *f* (*t*) = *f*_*Y X*_ (*t*) depend on both the pre- and postsynaptic populations *X* and *Y*, respetively. *V*_m_(*t*) denotes the postsynaptic membrane potential and ∈ {*E*_syn_ *E*_syn*E*_, *E*_syn*I*_} denotes the reversal potential of the synapse which is determined by the presynaptic cell type (i.e., excitatory or inhibitory). For simplicity, we will assume that 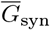 is independent of position. We will also assume that 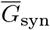 is static, that is, there are no synaptic plasticity rules or stochastic processes in place. Individual weights are, however, drawn from a distribution described by a probability density function 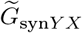, that is, 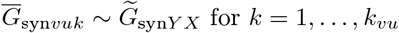. The subfix *vuk* in 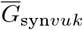 denotes the value for the *k*’th synapse between pre- and postsynaptic neurons *u* and *v*, respectively.
9. Let the conduction delays resulting from presynaptic action potential generation time to activation time of the synapse be greater than zero and randomly drawn from some distribution as 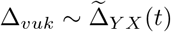. For simplicity, we let conduction delays be independent of cell location and geometry.
10. Let the sequence of spike times 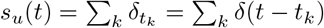 of each presynaptic neuron *u* in each population *X* is recorded throughout the entire simulation duration [0, *t*_sim_⟩. We choose to relax this requirement if a population in *X* represents an external population feeding persistent, uncorrelated events with spectrally ‘flat’ spiking statistics (e.g., fixed-rate Poisson point processes) into the recurrently connected network. We here (and for the remainder of this study) use the compact notation 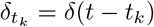 to denote Dirac delta functions centered around the time *t*_*k*_.
11. Let the weighted, directed graph representing edges (synaptic connections) between nodes (neurons) for every pair of pre- and postsynaptic populations be stored for purpose of validating the ‘hybrid scheme’ simulations described in the section Hybrid scheme for extracellular signal predictions. This storage requirement may also be relaxed if the total number of synapses over all connections is large enough to make storage infeasible or one could recreate the full connectivity graph procedurally (at least statistically). Graph weights represent maximum synaptic conductances. The graph also includes the synaptic locations on the postsynaptic neurons, and we will hereby let compartment index *m* equate to this location.
12. Let each postsynaptic neuron *v* in any population in *Y* be modeled using the ‘standard’ MC neuron formalism such that their transmembrane currents 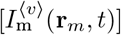 per compartment indexed by *m* can be computed. **r**_*m*_ denotes their midpoint coordinates.
13. Let extracellular signal contributions in different spatial locations (or axes in terms of current dipole moment) be computed and summed up as 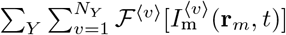. The matrix *ℱ*^⟨*v*⟩^ here denotes a linear mapping of transmembrane currents of cell *v* to a linearly dependent extracellular signal. The different forward models considered in this study are detailed in Volume-conductor forward models.

**Table 1.** MC neuron and recurrent network parameters (continued in Table 2).

| Symbol | Value/definition | Description |
| --- | --- | --- |
| $X$ | {Excitatory (E), inhibitory (I)} | Presynaptic cell types/populations |
| $Y$ | $X$ | Postsynaptic cell types/populations |
| $N_Y \in \{N_E, N_I\}$ | {8192, 1024} | Population sizes |
| $r$ | $\sqrt{x^2 + y^2}$ | Radius around vertical $z$-axis |
| $\mathcal{N}(\mu, \sigma)(x)$ | $\frac{1}{\sqrt{2\pi\sigma^2}} \exp -\frac{(x-\mu)^2}{2\sigma^2}$ | Gaussian distribution |
| $\tilde{\mathbf{r}}_X$ | $\mathcal{N}(0, 75\mu\text{m})(z) * r^2$ for $r \in [0, 150 \mu\text{m}]$ | Cell body probability density function |
| $C_{YX}$ | 0.05 for all $Y$ and $X$ | Pairwise connection probability<br>(Pairwise Bernoulli; no autapses) |
| $\tilde{k}_{YX}$ | $\begin{cases} \mathcal{N}(2, 0.5)(x_i), x_i \in \{1, 2, \dots, 20\}, \text{ for } X = \text{E} \\ \mathcal{N}(5, 1)(x_i), x_i \in \{1, 2, \dots, 20\}, \text{ for } X = \text{I} \end{cases}$ | Multapse probability mass function |
| $\mathcal{S}$ | {soma, apic, basal} | Morphology sections |
| $L_S$ | $\begin{cases} \{30, 1000, 200\} \mu\text{m} \text{ for } X = \text{E} \\ \{30, 200, 200\} \mu\text{m} \text{ for } X = \text{I} \end{cases}$ | Section length |
| $d_S$ | $\begin{cases} \{30, 3, 2\} \mu\text{m} \text{ for } X = \text{E} \\ \{15, 2, 2\} \mu\text{m} \text{ for } X = \text{I} \end{cases}$ | Section diameter |
| $n_{\text{seg}}^{\mathcal{S}}$ | $\begin{cases} \{1, 21, 5\} \text{ for } X = \text{E} \\ \{1, 5, 5\} \text{ for } X = \text{I} \end{cases}$ | # of segments per section |
| $c_m$ | $1 \mu\text{F cm}^{-2}$ | Membrane capacitance |
| $R_a$ | $100 \Omega \text{ cm}$ | Axial resistivity |
| $g_L^{\mathcal{S}}$ | {0.0000338, 0.0000589, 0.0000589} $\text{S cm}^{-2}$ | Passive leak conductance |
| $E_L$ | $-90 \text{ mV}$ | Passive leak reversal potential |
| $\bar{g}_{\text{Na}_t}^{\mathcal{S}}$ | {2.04, 0.0213, 0.0213} $\text{S cm}^{-2}$ | $\text{Na}_t$ conductance |
| $E_{\text{Na}}$ | $50 \text{ mV}$ | $\text{Na}^+$ reversal potential |
| $\bar{g}_{\text{Kv}_{3.1}}^{\mathcal{S}}$ | {0.693, 0.000261, 0.000261} $\text{S cm}^{-2}$ | $\text{Kv}_{3.1}$ conductance |
| $E_K$ | $-85 \text{ mV}$ | $\text{K}^+$ reversal potential |
| $\bar{g}_{\text{I}_h}^{\mathcal{S}}$ | {0.0002, 0.002, 0.002} $\text{S cm}^{-2}$ | $\text{I}_h$-current conductance |

The leak, fast inactivating $\text{Na}^+$ ( $\text{Na}_t$ ), fast, non-inactivating $\text{K}^+$ ( $\text{Kv}_{3.1}$ ) channel and non-specific cation current ( $\text{I}_h$ ) dynamics are those given in detail in [30]. Capped and discrete distributions are normalized such that the integral or sum over all values equals 1.

**Table 2.** Synaptic parameters for recurrent network (continued from Table 1).

| Symbol | Value/definition | Description |
| --- | --- | --- |
| $\tilde{G}_{\text{syn}YX}$ | $\begin{cases} \mathcal{N}(0.15 \text{ nS}, 0.02 \text{ nS})\Theta(G) & \text{for } X = \text{E}, Y = \text{E} \\ \mathcal{N}(0.125 \text{ nS}, 0.0125 \text{ nS})\Theta(G) & \text{for } X = \text{E}, Y = \text{I} \\ \mathcal{N}(4.5 \text{ nS}, 0.45 \text{ nS})\Theta(G) & \text{for } X = \text{I}, Y = \text{E} \\ \mathcal{N}(2.0 \text{ nS}, 0.2 \text{ nS})\Theta(G) & \text{for } X = \text{I}, Y = \text{I} \end{cases}$ | Synaptic conductance distributions |
| $E_{\text{syn}X}$ | 0 mV for $X = \text{E}$, -80 mV for $X = \text{I}$ | Synaptic reversal potential |
| $f_{YX}(t)$ | $\left( \frac{e^{-(t-t_s)/\tau_1} - e^{-(t-t_s)/\tau_2}}{e^{-\tau_{\text{peak}}/\tau_1} - e^{-\tau_{\text{peak}}/\tau_2}} \right) \Theta(t - t_s)$<br>where $\tau_{\text{peak}} = \frac{\tau_2 \tau_1}{\tau_2 - \tau_1} \log(\frac{\tau_2}{\tau_1})$ | Synaptic temporal kernel |
| $\tau_1$ | 0.2 ms for $X = \text{E}$, 0.1 ms for $X = \text{I}$ | Synaptic rise time constant |
| $\tau_2$ | 1.8 ms for $X = \text{E}$, 9.0 ms for $X = \text{I}$ | Synaptic decay time constant |
| $\tilde{\Delta}_{YX}$ | $\begin{cases} \mathcal{N}(1.5 \text{ ms}, 0.3 \text{ ms})\Theta(t - 0.3 \text{ ms}) & \text{for } X = \text{E}, Y = \text{E} \\ \mathcal{N}(1.4 \text{ ms}, 0.4 \text{ ms})\Theta(t - 0.3 \text{ ms}) & \text{for } X = \text{E}, Y = \text{I} \\ \mathcal{N}(1.3 \text{ ms}, 0.5 \text{ ms})\Theta(t - 0.3 \text{ ms}) & \text{for } X = \text{I}, Y = \text{E} \\ \mathcal{N}(1.2 \text{ ms}, 0.6 \text{ ms})\Theta(t - 0.3 \text{ ms}) & \text{for } X = \text{I}, Y = \text{I} \end{cases}$ | Conduction delay distributions |
| $L_{YX}(z)$ | $\begin{cases} \frac{\mathcal{N}(0,100)(z)}{3} + \frac{2\mathcal{N}(500,100)(z)}{3}, & X = \text{E}, Y = \text{E}, \mathcal{S} \setminus \{\text{soma}\} \\ \mathcal{N}(50, 100)(z), & X = \text{E}, Y = \text{I}, \mathcal{S} \setminus \{\text{soma}\} \\ \mathcal{N}(-50, 100)(z), & X = \text{I}, Y = \text{E} \\ \mathcal{N}(-100, 100)(z), & X = \text{I}, Y = \text{I} \end{cases}$ | Depth-dependence, syn. density |
| $\bar{G}_{\text{syn}Y\text{ext}}$ | 0.2 nS | External synapse conductance |
| $E_{\text{ext}}$ | 0 mV | Ext. synapse rev. potential |
| $f_{Y\text{ext}}(t)$ | $f_{YX}(t)$ | Ext. synapse temporal kernel |
| $\tau_1$ | 0.2 ms | Ext. synapse rise time constant |
| $\tau_2$ | 1.8 ms | Ext. synapse decay time constant |
| $k_{Y\text{ext}}$ | {465, 160} | # ext. synapses per neuron |
| $\langle \nu_{\text{ext}} \rangle$ | 40 s<sup>-1</sup> (Poisson statistics) | Ext. syn. activation rate |
| $\tilde{\Delta}_{Y\text{ext}}$ | $\delta(t)$ | Ext. syn. conduction delay |
| $L_{Y\text{ext}}$ | 1 | Ext. syn. depth dependence |

Capped and discrete distributions are normalized such that the integral or sum over all values equals 1.

**Table 3.**
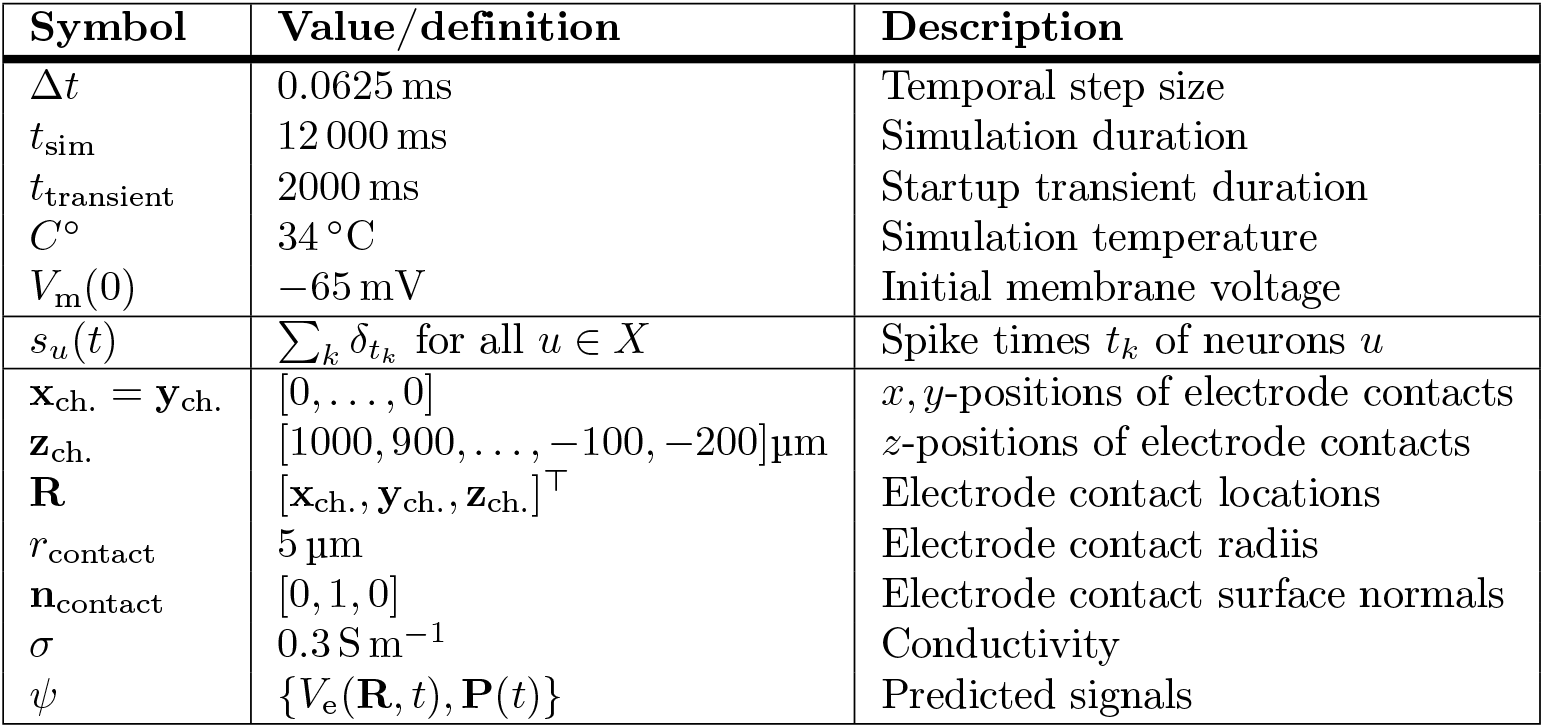
Measurement and simulation parameters for recurrent network (continued from Table 2).

| Symbol | Value/definition | Description |
| --- | --- | --- |
| $\Delta t$ | 0.0625 ms | Temporal step size |
| $t_{\text{sim}}$ | 12 000 ms | Simulation duration |
| $t_{\text{transient}}$ | 2000 ms | Startup transient duration |
| $C^\circ$ | 34 °C | Simulation temperature |
| $V_m(0)$ | -65 mV | Initial membrane voltage |
| $s_u(t)$ | $\sum_k \delta_{t_k}$ for all $u \in X$ | Spike times $t_k$ of neurons $u$ |
| $\mathbf{x}_{\text{ch.}} = \mathbf{y}_{\text{ch.}}$ | [0, ..., 0] | $x, y$-positions of electrode contacts |
| $\mathbf{z}_{\text{ch.}}$ | [1000, 900, ..., -100, -200] $\mu\text{m}$ | $z$-positions of electrode contacts |
| $\mathbf{R}$ | $[\mathbf{x}_{\text{ch.}}, \mathbf{y}_{\text{ch.}}, \mathbf{z}_{\text{ch.}}]^\top$ | Electrode contact locations |
| $r_{\text{contact}}$ | 5 $\mu\text{m}$ | Electrode contact radii |
| $\mathbf{n}_{\text{contact}}$ | [0, 1, 0] | Electrode contact surface normals |
| $\sigma$ | 0.3 S m<sup>-1</sup> | Conductivity |
| $\psi$ | $\{V_e(\mathbf{R}, t), \mathbf{P}(t)\}$ | Predicted signals |

### Reference networks of simplified ball-and-sticks neurons

The two-population, recurrent MC neuron network models constructed for this study, fully specified by the enumerated list above and parameter values listed in Table 1, 2 and 3, are kept intentionally simple for clarity of results. One main simplification is stylized neuron models with only a subset of ion channels distributed onto soma and dendritic compartments. The ‘E’ cell represents a phenomenological excitatory unit, while the ‘I’ cell represents a phenomenological inhibitory unit. Both share the same subset of passive and active ion channels taken from a biophysically detailed cell model [30], important for action potential generation (transient sodium, Na_t_; fast, non-activating potassium, SK_v1.3_) and sub-threshold dynamics (non-specific cation current, I_*h*_). Refer to [30] for details on these ion-channel dynamics. Network parameters were initially tuned by a combination of hand-tuning and parameter value scans, aiming to generate population spiking activity that is asynchronous and irregular (AI) [50] and with biologically plausible averaged spontaneous spike rates (approximately ⟨*v*_E_(*t*) ⟩ = 2.5 spikes s^−1^ for the ‘E’ population; ⟨*v*_I_(*t*) ⟩ = 5 spikes s^−1^ for the ‘I’ population).

### Reference networks of biophysically detailed neuron models

As an additional test of the methodology developed around the above description of a recurrent network of simplified ball-and-sticks MC neuron models, we replace the ‘excitatory’ (E) cell type with a biophysically detailed model of a thick-tufted layer 5b pyramidal cell of rat somatosensory cortex [30]. Here, we use model parameter values shown to produce acceptable BAC firing and perisomatic step current firing as summarized in [30, Table 3]. Each individual cell instance in the modified network model is rotated by 4.729 rad and −3.166 rad around the horizontal *x*− and *y*−axes, respectively, in order to first align the apical dendrite with the vertical *z*−axis, before applying a random rotation around the *z*−axis. By increasing the number of extrinsic synapses distributed on each neuron to 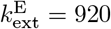, the typical population firing rates, and network state is well preserved when compared to the reference network. All other parameters remain as defined in Tables 1 to 3.

### Reference neuron networks with perturbed synaptic conductances

Parts of this study are devoted to the effect of perturbed network parameters in different network instances on our proposed methodology. For this, we incorporated a connection weight scaling factor *J* ∈ {0.975, 1, 1.025, 1.05, 1.075}, and rescaled recurrent synaptic max conductances 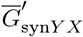 (and parameters derived from them) as

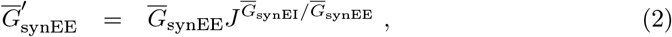

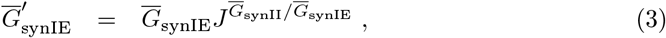

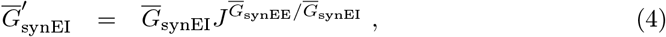

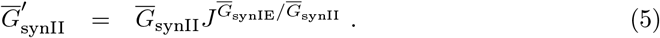

Effectively, perturbing *J* shifts the relative balance of excitatory and inhibitory synaptic input in the networks. A factor *J* = 1 corresponds to our unperturbed reference network.

### Leaky integrate-and-fire (LIF) point-neuron network

As a proof of principle that the ‘kernel method’ (see Kernel-based extracellular signal predictions) can be utilized for ‘live’ extracellular signal predictions with spiking point-neuron network models or other types of networks with abstract neuron representations, we fit connectivity parameters of a phenomenological two-population network of leaky integrate-and-fire (LIF) point neuron network with current-based synapses to mimic the spiking activity of the unperturbed reference network of ball-and-sticks neurons. After initial hand tuning of the network parameters into a reasonable state of activity resembling the reference network’s state, we subsequently used the multi-objective optimization NSGA-II non-dominated sorting genetic algorithm [51] in order to fine tune key network connectivity parameters, namely synaptic weights *Ī*_syn*Y X*_, membrane capacitance of neurons in each population *C*_m*X*_, weight of extrinsic synapses 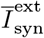, and mean value of the conduction delay distributions 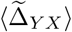. The full network and neuron descriptions are given in Table 4, including best-fit parameters and parameter value boundaries used for the fitting procedure. The network is implemented and simulated in NEST [52, 53], using exact integration for step size Δ*t* [54].

**Table 4.**
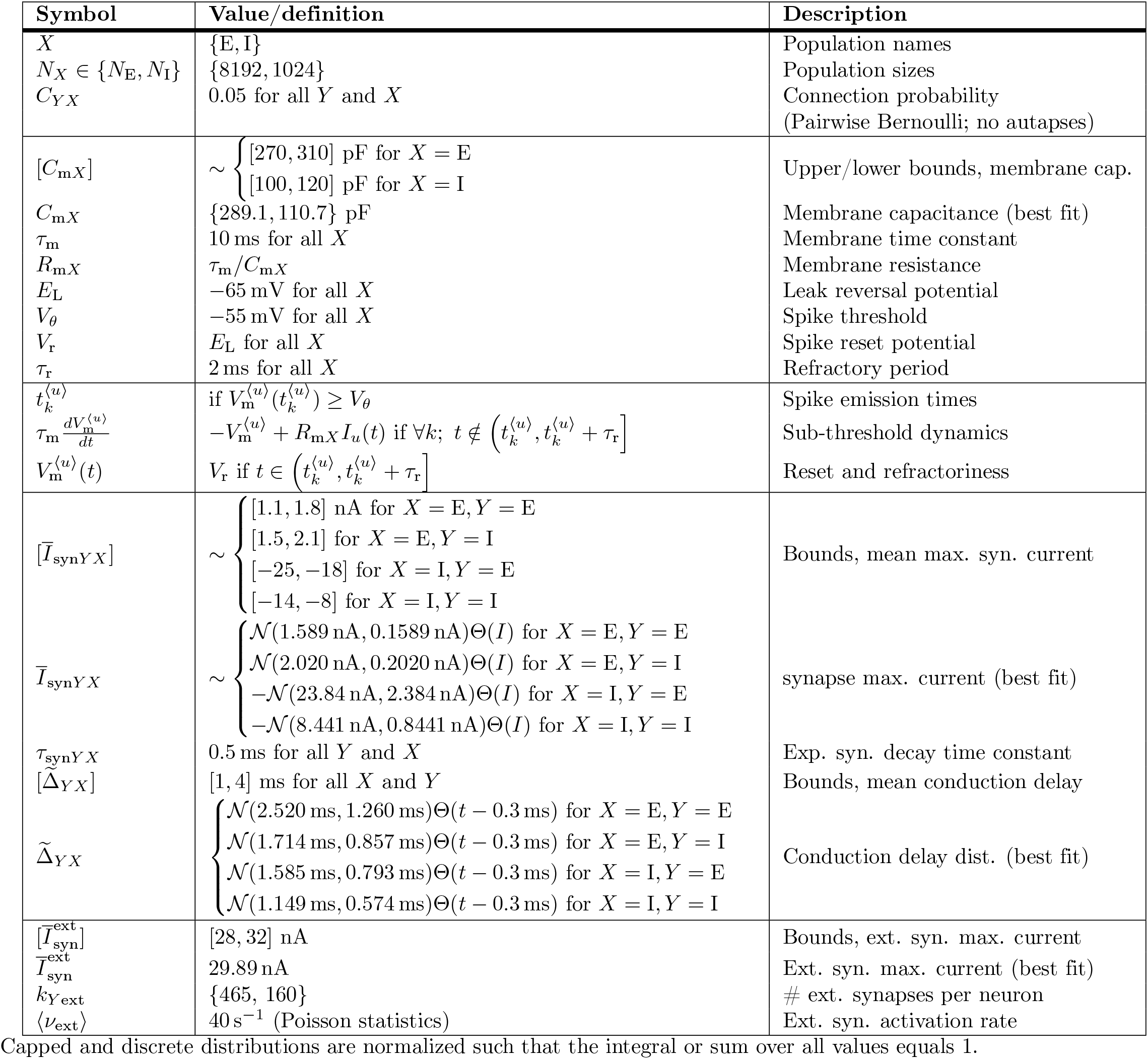
LIF network and neuron parameters.

For the parameter fitting, we used the implementation of the NSGA-II class `pymoo.algorithms.nsga2.NSGA2` provided by the `pymoo` Python package [55]. We defined the objective functions to be minimized using the `pymoo.optimize.minimize` method as

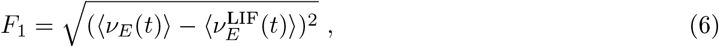

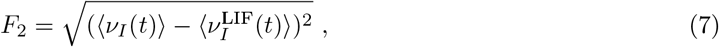

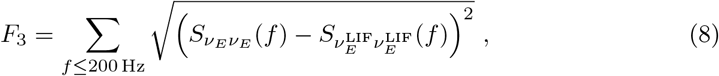

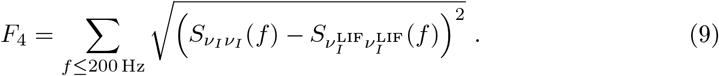

Here, *v*_*X*_ (*t*) and 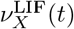 denote population spike rates of the MC and LIF neuron network populations, respectively. 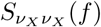 and 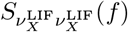 denotes population spike rate autospectral power at each frequency *f* (see Signals and signal analysis methods for details). For this minimization problem, we used an initial population size of 100, and ran the algorithm for 20 generations with default parameters.

The pseudo-weight vector approach [55, 56] is used to select a solution from the solution set that performs well with respect to all objective functions. The pseudo weight, a normalized distance measure from the worst solution for each objective function *F*_*i*_, is herein calculated as:

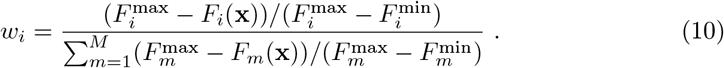

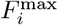 and 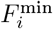 denotes the maximum and minimum value of *F*_*i*_(**x**) in the last generation, respectively. Then, the best-fit parameter vector **x** where chosen as the one that minimized ∥ [*w*_1_, *w*_2_, *w*_3_, *w*_4_]^⊤^ − [0.25, 0.25, 0.25, 0.25]^⊤^ ∥. We here use the implementation provided by the `pymoo.factory.get_decision_making` method.

### Hybrid scheme for extracellular signal predictions

The so-called ‘hybrid scheme’ [18] is a proposed solution for computing extracellular signals from spiking activity in recurrent neuron network models. This scheme is hybrid in the sense that the spiking activity of recurrent networks is first simulated separately and stored, then stored spike events are loaded and used for synaptic activation times in corresponding populations of MC neuron models set up to predict extracellular signals. In this latter step, synapses are placed on postsynaptic neurons and are activated at times as they would occur in the corresponding recurrent network, negating recurrent connections and spike communication between MC neurons. Thus, the problem of computing signals can be solved in an embarrassingly parallel manner. As the scheme relies on prerecorded spike events, our application of the scheme employs postsynaptic neurons that do not generate APs.

Here, we incorporate the hybrid scheme by storing population geometries, spikes, and the full synaptic connectome (placements, weights, conduction delays, pre- and postsynaptic neuron IDs) of the recurrent MC neuron networks to file, and reinstate synaptic placements and activation times in separate simulations without actual recurrent connections. Locations and activation times of extrinsic synapses are not stored directly due to their large count. Here we ensured replicable placements and activation times by fixing the random seeds affecting these. This step allows for computing signals identical to the recurrent model in case the MC neuron models are those of the recurrent network, but here, we shall rely on models where the membrane and synapse dynamics are approximated by linear dynamics. Thus, only signal contributions that stem from synapse activations on postsynaptic neurons are accounted for in predicted signals, while contributions by presynaptic APs are not. The scheme thus lends itself to predictions of signals thought to mainly stem from synaptic activity, that is, LFP, ECoG, EEG, and MEG signals.

### Linear approximations to synapse and membrane dynamics

Here, we describe the different *linear* approximations to the different constituents of the conductance-based *non-linear* recurrent neuron network models, calculated via the following steps:

1. Approximate conductance-based synapses by equivalent current-based synapses:

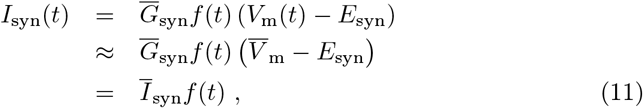

where 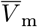 denotes the typical postsynaptic potential (or its expectation value). The synapse current magnitude *Ī* _syn_ is constant. Here, we typically recorded somatic potentials in a subset of neurons in each population of the recurrent MC neuron networks and let 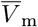 equate the median somatic potentials. Where noted, we perturb 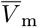 by some value or use values obtained across the neuronal morphologies.
2. Active ion channel currents on the specific form

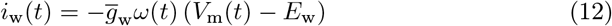

are approximated by equivalent, linearly dependent currents similar to [40, 57]. Here, *E*_w_ is the channel reversal potential and *ω*(*t*) the gating variable which dynamics are given in terms of an activation time function *τ*_w_(*V*) and activation function *ω*_∞_(*V*) as

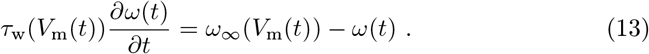 If the voltage dynamics of the active compartment is defined by

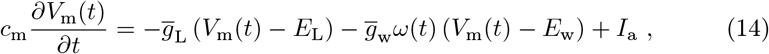

one can obtain the so-called *quasi-active approximation* [40, 57, 58] by linearizing each voltage dependent term around the steady state value *V* _m_ resulting in

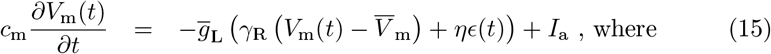

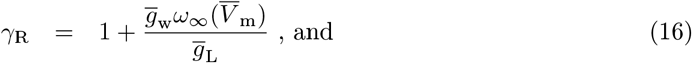

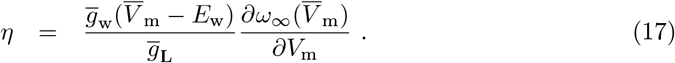 Here, an equivalent gating variable is defined as

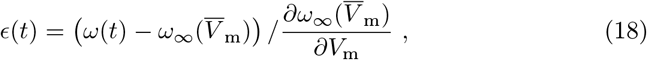

and its linear dynamics is governed by

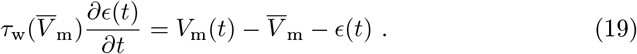 Above, *γ*_R_ denotes the ratio between the total and leak conductance, while *η* characterize whether the quasi-active current approximation acts as positive (*η <* 0) or negative (*η >* 0) feedback. For the special case *η* = 0 the quasi-active current is ‘frozen’, acting as a passive current [40]. Note that the above sets of equations correspond to channels usually modeled with a single state variable (e.g., I_h_-type currents), but generalize also to current types with more than one gating variable (e.g., Na^+^- and K^+^ type currents), see [57] for details.
3. With linearized active ion channels, the leak reversal potential *E*_L_ is further modified as

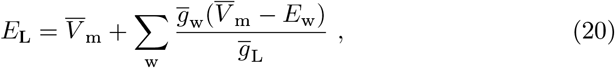

which ensures that the resting potential of the quasi-active model is similar to 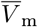. We note that this modification do not affect extracellular signal predictions where current-based synapses (pt. 1) are used, but is applied anyway as in [40].
4. In principle one may remove active ion channels omitting the above linearization tricks altogether if their net contributions to the total transmembrane currents can be assumed to be minuscular around typical membrane voltage values. Here however we do account for all channels.
5. With current-based synapses (pt. 1), we optionally incorporate the *effective membrane conductance g*_eff_ which amounts to a modified passive leak conductivity. Assuming the total membrane conductance depends only on synaptic currents of recurrent and external connections and the passive leakage current, the total leak membrane conductivity per postsynaptic compartment *m* of postsynaptic neuron indexed by *v* is

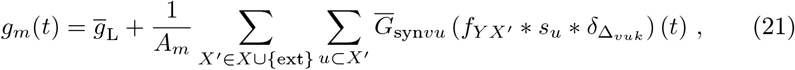

where 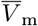 is the specific passive leak conductance, *A*_*m*_ compartment area, *s*_*u*_(*t*) the sequence of presynaptic spikes and 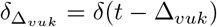 the conduction delay. The asterisk symbol (∗) denotes a temporal convolution. The double sum over presynaptic populations *X*^′^ ∈ *X* ∪ {ext} and units *u* ∈ *X*^′^ implies that each presynaptic unit *u* targeting the compartment is accounted for. We introduce this notation to express that also synapses from external sources (‘ext’) must be accounted for. Assuming a fixed average presynaptic spike rate ⟨*v*_*u*_(*t*)⟩ and a normalized delay distribution (where 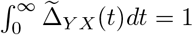), the time-averaged effective conductance in each compartment *m* is approximately

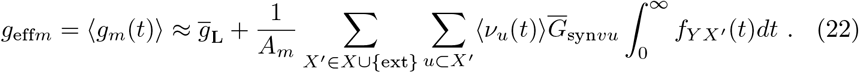 Then, the original 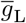 value may be replaced by *g*_eff*m*_ on a per-compartment basis. Note that we compute values of *g*_eff*m*_ independently from contributions by linearized active ion-channel contributions (pt. 2), which still contribute to the total sum of conductances.

### Kernel-based extracellular signal predictions

In case the relations between spikes in presynaptic populations and resulting extracellular signals arising mainly from evoked responses in targeted postsynaptic populations are approximately linear, filter- or ‘kernel’-based prediction methods may greatly simplify signal predictions at the level of populations. If we first define the presynaptic population spiking activity as the sequence of Dirac delta functions *s*_*X*_ (*t*) = Σ_*u∈ X*_ *s*_*u*_(*t*), the signal approximation 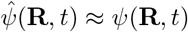 may be computed as the sum over linear convolutions

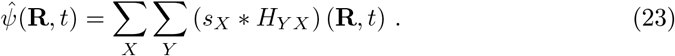

Here, *H*_*Y X*_ (**R**, *τ*) are representative spatiotemporal spike-signal impulse responses for pairs of pre- and postsynaptic populations measured relative to presynaptic spike events at time lag *τ* = 0. As we deal with spike events and sampled signals on a discrete time grid, it is convenient to redefine the spike sequences *s*_*X*_ (*t*) as spike rates by the temporal binning

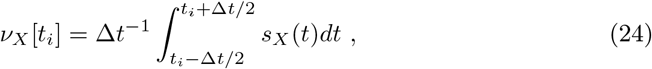

where Δ*t* denotes the simulation step size. Then, the above equation can be written as

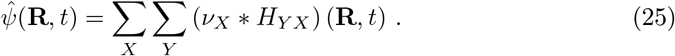

Throughout this section, we describe two alternative methods to compute such kernels *H*_*Y X*_ (**R**, *τ*), either via the hybrid scheme as in [18], or directly as described below. As the kernels are equivalent to finite impulse responses (FIR) as they are 0 for all time lags *τ <* Δ*t*, the linear convolutions can be replaced by a linear filter function implementation (see Signals and signal analysis methods below).

#### Kernel predictions via the hybrid scheme

The linear cable equation combined with linearized synapse- and ion-channel dynamics in our application of the hybrid scheme (cf. Hybrid scheme for extracellular signal predictions) provides a relatively straightforward method to compute representative sets of spatiotemporal kernel functions as in [18]. This earlier study shows that a good approximation to the signal *ψ*(**R**, *t*) can be obtained by first measuring kernels averaged over all pre- and postsynaptic neurons in each population *X* and *Y*. In order to compute these kernel averages directly using the hybrid scheme, actual network spiking activity is first replaced by simultaneous and deterministic events 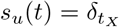 where *t*_*X*_ *>* 0 is a chosen time for each population *X*, then the signal contributions of each postsynaptic population *ψ*_*Y X*_ (**R**, *t*) is computed via the disassociated network model around *t*_*X*_; and the response is averaged over the presynaptic neurons as

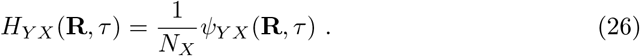

Here, *τ* denote time relative to *t*_*X*_. Thus these kernels must be causal, that is, by construction *H*_*Y X*_ (**R**, *τ*) = 0 for *τ <* 0 as any contribution to the signal *ψ*(**R**, *t*) is solely postsynaptic. No signal contributions before the presynaptic spike event at *τ* = 0 plus the minimum conduction delay is accounted for^1^. We let the computed kernels span the interval *τ* ∈ [−*τ*_max_, *τ*_max_], where *τ*_max_ denotes a maximum lag value. The postsynaptic responses typically rise and decay back to approximately zero after a few tens of milliseconds. This decay time is related to the time constants relevant to the neuronal dynamics (that is, *τ*_m_, *τ*_syn_, *τ*_w_, …). Throughout this manuscript, we chose *τ*_max_ = 100 ms for computed kernels, which we assume is a few multiples of relevant time constants.

#### Direct kernel predictions from single MC simulations

Different from the hybrid scheme kernel method described above, the main aim of this work is to develop a method to directly compute a set of accurate and deterministic kernels *Ĥ*_*Y X*_ (**R**, *τ*) needed for all connection pathways between pre- and postsynaptic populations *X* and *Y*, based on some expectation values for cell and synaptic placements and other network parameters. We aim to replace simulations of populations of MC neurons via the hybrid scheme with a single MC neuron simulation per kernel. Thus the number of MC neuron simulations corresponds to the number of pathways between any population *X* and *Y* which is significantly less than the total neuron count in each network. The hat denotes kernels computed using this direct method, in contrast to hybrid scheme kernels. First, we assume that:

1. The dynamics of the neuronal cables and synaptic input can be approximated as a linear system resulting from the same steps as in Linear approximations to synapse and membrane dynamics.
2. Each postsynaptic population can be represented by *one* typical biophysically detailed neuron model. Effectively, the whole postsynaptic neuron population is collapsed to a single neuron with linearized membranes receiving all inputs, while the effect of the spatial distribution of cells in space is accounted for via the VC forward model (see Modified forward models for deterministic kernel predictions).
3. The underlying statistics of synaptic placements and currents are preserved, which allows us to compute the average synaptic current density for each recurrent connection over the whole postsynaptic population ‘neuron’. Accounting for the distribution of neurons along the *z*-axis and ignoring their radial location, we let the synaptic density be proportional to the membrane area of postsynaptic compartments *A*_*m*_ multiplied by a function *ℒ*_*Y X*_ (*z*) obtained as the convolution of *L*_*X Y*_ (*z*) (see Table 2) and the *z*−component of 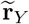 (defined in Table 1). Hence we compute the expectation value for synaptic in-degree per compartment indexed by *m* as

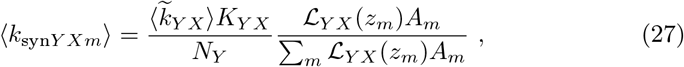

where *z*_*m*_ denotes the midpoint location of each compartment projected on the *z*-axis, and 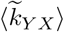 the mean multapse count per connection. With this quantity one may define the per-compartment synaptic input per activation as

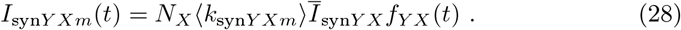 As above, the term *f*_*Y X*_ (*t*) denotes the temporal component of synapse currents for each connection.
4. Optionally accounting for the effective leak conductivity, Eq 22 must be modified per compartment as

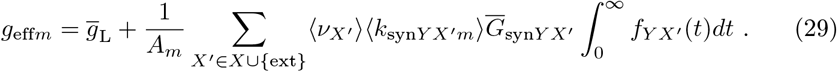 As above, we account for the external population ‘ext’ jointly with the main network populations in *X*.
5. Then one may straightforwardly compute the resulting postsynaptic response, that is, the full set of transmembrane currents [*I*_m_(**r**_*m*_, *τ*)], by applying synaptic currents *I*_syn*Y Xm*_(*τ*) in a single MC neuron simulation for all connections between populations *X* and *Y*. In order to temporarily compute the approximated kernel functions 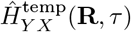 for different extracellular signals, the resulting transmembrane currents must be combined with appropriate forward model matrices *ℱ* calculated as described below.
6. Finally to account for network conduction delay distributions, the intermediate kernels must be filtered in the temporal domain as

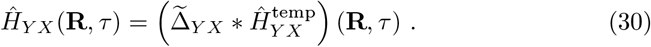

### Volume-conductor forward models

#### Forward models for reference-network and hybrid scheme signals

As derived from *volume conductor theory*, the different electric and magnetic signals that can be computed from the electric activity of brain cells are *linearly* dependent on transmembrane currents (see e.g., [5] and references therein). Thus, some arbitrary signals *ψ*(**R**_*n*_, *t*) in *M* different spatial locations **R**_*n*_ (or directions in case of current dipole moments) from *N* compartmental sources indexed by *m* located at **r**_*m*_ can be computed as

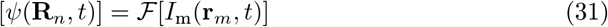

where *ℱ* is a matrix with dimensions (*M, N*) wherein each element *f*_*nm*_ is the chosen forward solution mapping the contribution from each source to the corresponding measurement. [*I*_m_(**r**_*m*_, *t*)] denotes the transmembrane currents of compartments *m* at time *t*. For the presently used line sources [11, Eq. (4)], the elements of *ℱ* are calculated using

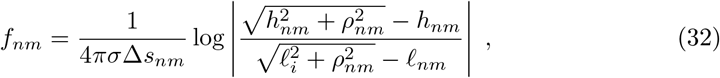

where *ρ*_*nm*_ is the distance perpendicular to line source (compartment) *m, h*_*nm*_ the longitudinal distance from the end of the line source and *ℓ*_*nm*_ = Δ*s*_*nm*_ + *h*_*nm*_ the longitudinal distance from the start of the line source with length Δ*s*_*nm*_ to some electrode contact located at **R**_*n*_. The line-source approximation assumes an infinite homogeneous, isotropic, and linear volume conductor with conductivity *σ*. Measurement sites are treated as infinitesimally small points, so to mimic the finite extent of contacts of experimental recording electrodes, we apply the ‘disk-electrode’ approximation to the extracellular potential [11, Eq. (6)] by embedding averaged values of *f*_*nm*_ from Eq 32 for 100 random locations within radius *r*_contact_ into *ℱ*.

The approach applies also to other types of measurements that are linearly dependent on the transmembrane current sources, such as the current dipole moment [39]. For calculations of the current dipole moment **P** the columns of *ℱ* are simply

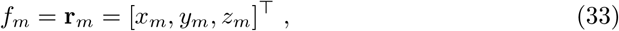

where (*x*_*m*_, *y*_*m*_, *z*_*m*_) denotes the midpoint coordinates of each compartment.

#### Modified forward models for deterministic kernel predictions

When computing extracellular signals via the kernel predicting scheme we must account for the distributions of cells in space. Here we assume that each population is radially symmetric around the vertical *z*-axis, homogeneous within some radius *R* and inhomogeneous along the *z*−axis as described by a probability density function 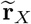 (see Table 1). In order to compute extracellular potentials, we use the analytical forward solution for the electric potential from a planar disk with homogeneous current density [59]

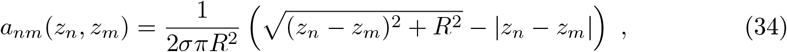

which is subsequently convolved with the depth-dependence of cell placement 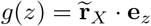 (where **e**_*z*_ denotes the unit vector along the *z*−axis), resulting in matrix elements:

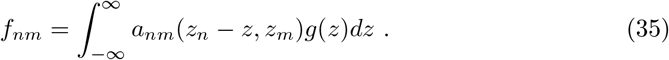

Here, we solve this convolution integral numerically using the `quad` method of the `scipy.integrate` module. Note that we apply the same equation also when predicting kernels for the biophysically detailed neuron network (see Reference networks of biophysically detailed neuron models). This formalism also assumes that the spread of 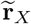 is ‘vast’ versus typical compartment lengths and contact radii. The planar disk radius *R* is set equal to the population radius *r*.

To compute the current dipole moment assuming radial symmetry around the *z*-axis the mapping matrix’ columns are simply modified as

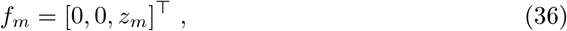

where *z*_*m*_ denotes the midpoint coordinates of each compartment along the *z*−axis. We do not account for the distribution of cells along the *z*-axis as it does not affect the current dipole moment. Due to radial symmetry, the components in the lateral directions are expected to cancel [12, 60], hence the corresponding matrix elements are set to zero.

### Signals and signal analysis methods

Throughout this study, the different signals we consider are: membrane potentials *V*_m_(*t*); spike trains *s*_*u*_(*t*); population firing rates *v*_*X*_ (*t*) obtained by counting spikes per time bin of width Δ*t* divided by bin width providing a signal with unit spikes s^−1^ as defined in Eq 24; and raw and low-pass filtered extracellular signals *ψ*(**R**_*n*_, *t*) (extracellular potentials *V*_e_(**R**_*n*_, *t*); current dipole moments **P**(*t*)). For extracellular signals we consider only frequencies *f >* 0Hz by subtracting the mean value in each channel for times *t > t*_transient_.

For low-pass filter operations, we used an elliptic (Cauer) digital filter design. Here, we used filters of the 2nd order with 0.1 dB maximum ripple in the passband, minimum attenuation of 40 dB in the stopband, and a critical (cutoff) frequency of 100 Hz. Filter coefficients were generated using the `scipy.signal.ellip` function with parameter `output=‘sos’` (second-order sections). The low-pass filter was applied to the data using the `scipy.signal.sosfiltfilt` function which implements a forward-backward (zero time-lag) filter operation.

In order to quantify relative differences in amplitudes of approximated signals *x*(*t*) and ground truth *y*(*t*) we defined the ‘ratio of standard deviations’ as

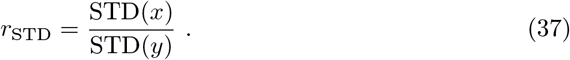

In order to quantify temporal agreement with signals *x*(*t*) and *y*(*t*) we computed the squared correlation coefficient (coefficient of determination) *R*^2^ at zero time lag as

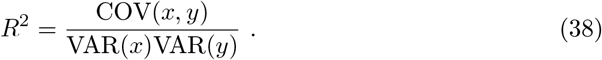

In order to aggregate our *R*^2^ and *r*_STD_ metrics for signals computed at different depths, we computed the 10th and 90th percentiles using the implementation of `numpy.quantile` with quantiles equal to 0.1 and 0.9, respectively.

For convolutions, we use the discrete convolution between vectors *x* and *y* defined as

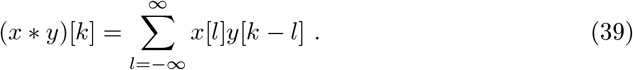

Here, we used the implementation provided by `numpy.convolve` with `mode=‘same’`.

Application of discrete FIR filter coefficients *h* to a signal *x* (relevant for NEST predictions) is defined as

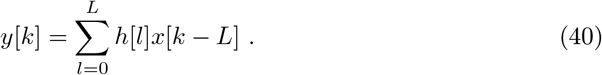

Estimates of cross power spectral densities (CPSD) *S*_*xy*_(*f*) and power spectral densities (PSD) *S*_*xx*_(*f*) of signals *x*(*t*) and *y*(*t*) use the Welch’s average periodogram method [61] as implemented by `scipy.signal.csd`. Unless specified otherwise, we use the periodogram settings `nfft=2048`, `noverlap=1536`, `fs=`Δ*t*^−1^ (in Hz) and `detrend=False`. When optimizing point-neuron network parameters we used the setting `detrend=‘constant’` when computing the features *F*_3_ and *F*_4_.

The real-valued coherence (magnitude-squared coherence) between signals *x*(*t*) and *y*(*t*) we compute via their CPSD and PSD functions as

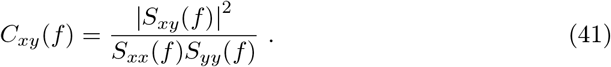

### NESTML FIR filter extension

In order to incorporate extracellular signal predictions using the computed sets of causal kernels from a point-neuron simulation in NEST (see Leaky integrate-and-fire (LIF) point-neuron network), a finite impulse response (FIR) filter implementation of Eq 40 is now expressed in the NESTML modeling language [62, 63]. The FIR filter model is written as a neuron model in NESTML, which takes neuronal spikes as input and computes the filter output while the simulation progresses. The output can then be queried and recorded to file using standard NEST devices. The NESTML toolchain generates C++ code for the model, which is compiled into a NEST extension module, allowing the FIR filter node (or a heterogeneous population of filter nodes) to be instantiated in NEST simulations.

As per Eq 40, the FIR filter model defines *L* as the order of the filter and *h* as a vector of length *L* containing the filter coefficients. The values of *L* and *h* can be set externally from the simulation script, and in this study we insert filter coefficients from each set of predicted kernels *Ĥ*_*Y X*_ (**R**_*n*_, *τ* ≥ 0) for each different extracellular signal (see Direct kernel predictions from single MC simulations for details). The input spikes are binned per time step, and the spikes for the last *L* time steps are stored in a circular buffer *x* of length *L*. At every time step during the simulation, the binned input spikes in *x* are multiplied with filter coefficients in vector *h* and summed according to Eq 40. The index to vector *x* is also adjusted such that the appropriate element of the circular buffer is accessed. The resulting filter output is stored in a (scalar) state variable, *y*, which can be recorded using a multimeter in NEST.

### Data availability and replicability

#### Codes and software tools

This study has been made possible using the following software tools: GCC 11.2.0, mpich 3.4.2, Python 3.9.6, ipython 7.27.0, jupyter-notebook 6.0.3, numpy 1.21.3, scipy 1.7.1, matplotlib 3.4.3, pandas 1.3.4, seaborn 0.11.2, pymoo 0.4.2.2, mpi4py 3.1.3, h5py 3.5.0, NEURON 8.0.2, MEAutility 1.5.0, LFPykit 0.4, LFPy 2.2.6, LFPykernels 0.1.rc8 (github.com/LFPy/LFPykernels, git SHA: 4fd79ab), NEST 3.1 (github.com/nest/nest-simulator, git SHA: 512022e54), NESTML 4.0-post-dev (github.com/nest/nestml, git SHA: 0b251ec), parameters 0.2.1 (github.com/NeuralEnsemble/parameters, git SHA:b95bac2).

In order to ensure *Methods and results reproducibility* [64, 65], all simulation codes required to replicate the findings reported here are publicly available at github.com/LFPy/LFPykernels. These include the reference implementation of the methodology which is installable via the usual Python distribution channels as:

- `pip install --pre lfpykernels` # or
- pip install git+https://github.com/LFPy/LFPykernels

The code repository also includes a Docker recipe file which may be used to build containers with the full software environment required by the simulations and analysis. Versioned releases of the `LFPykernels` tool is permanently deposited on Zenodo.org [66].

#### Hardware details

All computationally demanding simulations for recurrent networks and reconstructed networks of MC neurons as well as parameter optimizations were performed on the standard compute nodes of the JUSUF compute cluster at the Jülich Supercomputing Centre (JSC), Jülich Research Centre, Jülich, Germany. Each compute node has two AMD EPYC 7742 CPUs (2 × 64 physical cores) running at 2.25 GHz, 256 GB of DDR4 RAM running at 3200 MHz. The compute nodes are interconnected by InfiniBand HDR100 (Connect-X6). Each MC network simulation ran in parallel distributed across 8 compute nodes with 1024 Message Passing Interface (MPI) processes, using the ParTec ParaStation MPI implementation. Point-neuron network simulations were executed using 32 OpenMP threads, 1 core per thread. All relevant software tools were compiled with compilers from GCC.

Post-processing, calculations of deterministic kernels, other analysis, and plotting were performed on a MacBook Pro (13-inch, M1, 2020) with 16 GB RAM running macOS Big Sur (v11.6) with the Conda (conda.io) package management system with packages from the conda-forge channel (conda-forge.org).

## Results

### Neuron models with linearized membrane dynamics

The results presented throughout this study rely on three different fully active multicompartment (MC) neuron models, and versions where their voltage-dependent ion channel dynamics are linearized around a chosen membrane voltage value. These linearization steps are detailed in (pts. 2-3 under Linear approximations to synapse and membrane dynamics). The cell morphologies are shown in Fig 2A. The phenomenological ‘ball-and-sticks’ models ‘E’ and ‘I’ represent excitatory and inhibitory neurons in the two-population recurrent network in the following sections, while the biophysically detailed layer 5 pyramidal cell model ‘E_Hay2011_’ [30] later on replaces the ball-and-sticks ‘E’ population (in Methods performance using biophysically detailed cell models). The ‘E’ and ‘I’ neurons are both modeled with a single compartment for the soma, and dendritic sections pointing upwards and downwards along the depth axis.The ‘E’ cell has a prominent apical section 1 mm in length while the ‘I’ cell dendritic sections are symmetric around the soma.

**Fig 2.**
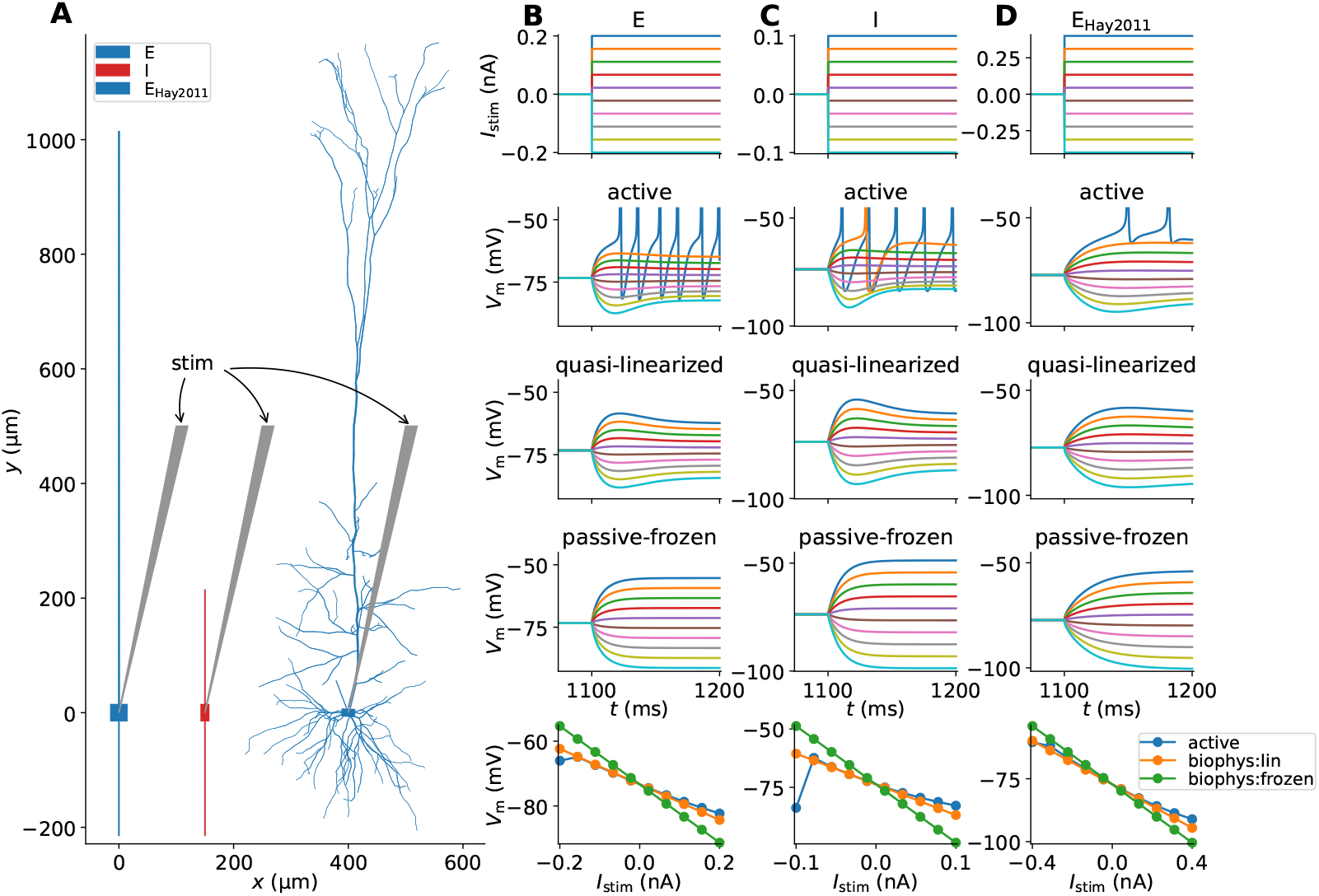
Model neurons and somatic responses with active and linearized ion-channel dynamics. (**A**) Neuronal geometries of neurons representing excitatory (E) and inhibitory (I) neurons, as well as a biophysically detailed pyramidal cell model (E_Hay2011_ [30]) replacing population ‘E’ in the modified network. (**B**) Excitatory (E) neuron responses in active and linearized versions. Row 1: Step input current with variable magnitude injected into the neuron’s soma compartment. Row 2: Somatic voltage responses to step input currents for the active neuron version. Colors corresponds to each respective trace in row 1. Row 3: Input responses in the quasi-active linearized (`biophys:lin`) version. Row 4: Input responses in the passive-frozen (`biophys:frozen`) version. Row 5: Response amplitudes at *t* = 1200 ms as function of stimulus magnitude. (**C**) Same as column B but for the inhibitory (I) neuron model. (**D**) Same as column B but for the biophysically detailed excitatory (E_Hay2011_) neuron model.

As a first check comparing active and linearized neuron dynamics in absence of synapses, we stimulate the different cell models with small step-like hyper- and depolarizing input currents to the somatic compartment. The dynamics are linearized around the steady state somatic membrane potentials in absence of stimuli. In panels B-C we compare responses of the ‘E’ and ‘I’ model versions with ‘quasi-active’ linearized versions of the I_h_-type channel (`biophys:lin`) plus frozen dynamics for the Na_t_- and SK_v1.3_ channels as well as the version were also the I_h_-type channel is frozen (`biophys:frozen`). In both cases, the quasi-active versions can capture the sub-threshold dynamics, including the sag and rebound effects explained by the I_h_-channel currents. The fully passive-frozen models are effectively similar to models with only passive leak channels, which is reflected in the corresponding responses. These models then perform worse with respect to capturing the sub-threshold dynamics of the fully active versions. Note also that these linearized model neurons can not generate APs for stronger depolarizing input currents, unlike their active counterparts. The row 5 panels show the response amplitudes after 100 ms stimulus duration, and unsurprisingly the linearized neuron dynamics are linearly dependent on stimulus amplitude. Quasi-active models match the corresponding active model responses well for small stimulus amplitudes.

In Fig 2D, the same experiment is performed with the biophysically detailed model neuron [30]. Here, a quasi-active version of the Na_P_ conductance is incorporated in addition to the quasi-active I_h_ channel, while remaining channels are in their passive-frozen states (`biophys:lin`). Again, the sub-threshold dynamics for small perturbations are captured by the quasi-active model in an excellent manner, resulting in similar responses below the firing threshold. Similar to our earlier observation, the model version with all passive-frozen dynamics (`biophys:frozen`) can not capture the somatic response accurately. The same qualitative observations hold true in case current input is delivered to a dendritic location approximately 200 µm from the soma (S1 Fig).

### Reference MC neuron network with extracellular signal predictions

Representing our reference networks for generating ground-truth extracellular signals, and spiking activity used for signal approximations, Fig 3A shows the populations of ball-and-sticks neurons and extracellular recording geometry for a phenomenological two-population MC neuron network set up according to pts. 1-13 in Reference multicompartment neuron networks. For this network (as well as networks with perturbed parameters), we predict extracellular potentials at depths highlighted by black circular markers treating compartments as line sources (Eq 32), as well as the current dipole moment (Eq 33). The current dipole moment determines EEG and MEG-like signals, as both can be computed from it using the appropriate forward model [3, 4, 12, 60]. Panels B and C show the distributions across depth of somas and instantiated synapses for each pair of pre- and postsynaptic populations, accordingly. All neurons receive depolarizing input by randomly distributed excitatory synaptic input with random activation times. A few somatic membrane potential traces recorded in each population is shown in panel D. The median values 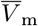 for a sample size of *N* = 1024 in each population are used for linearization of ion-channel and synapse dynamics in the following sections. The spike raster plot (Fig 3E) shows the resulting activity to be stable and asynchronous-irregular at biologically plausible rates. The I cells fire more often than the E cells on average, around 5.1 Hz and 2.6 Hz respectively. Oscillations at the level of the populations are clearly visible in the corresponding spike-count histograms (panel F) and rate spectra (panel G). These oscillations around 55 Hz can be expected to be expressed in extracellular signals, and indeed the extracellular potential (panel H) shows oscillations with varying amplitudes across depth. We note in passing that the generated extracellular potentials are in line with experimentally observed signals with amplitudes of a few 100 µV, with few visible extracellular spike signatures. The oscillations generated by the network are prominently captured also in the current dipole moment (panel I), however only in the vertical *z*−component *P*_*z*_. Due to the symmetry of the neural populations around the *z*−axis and the cell alignments along the same axis, the orthogonal components *P*_*x*_ and *P*_*y*_ cancel. Next, we investigate how these signals may be captured by models that only use MC neurons with linearized ion-channel and synapse dynamics and no recurrent connections.

**Fig 3.**
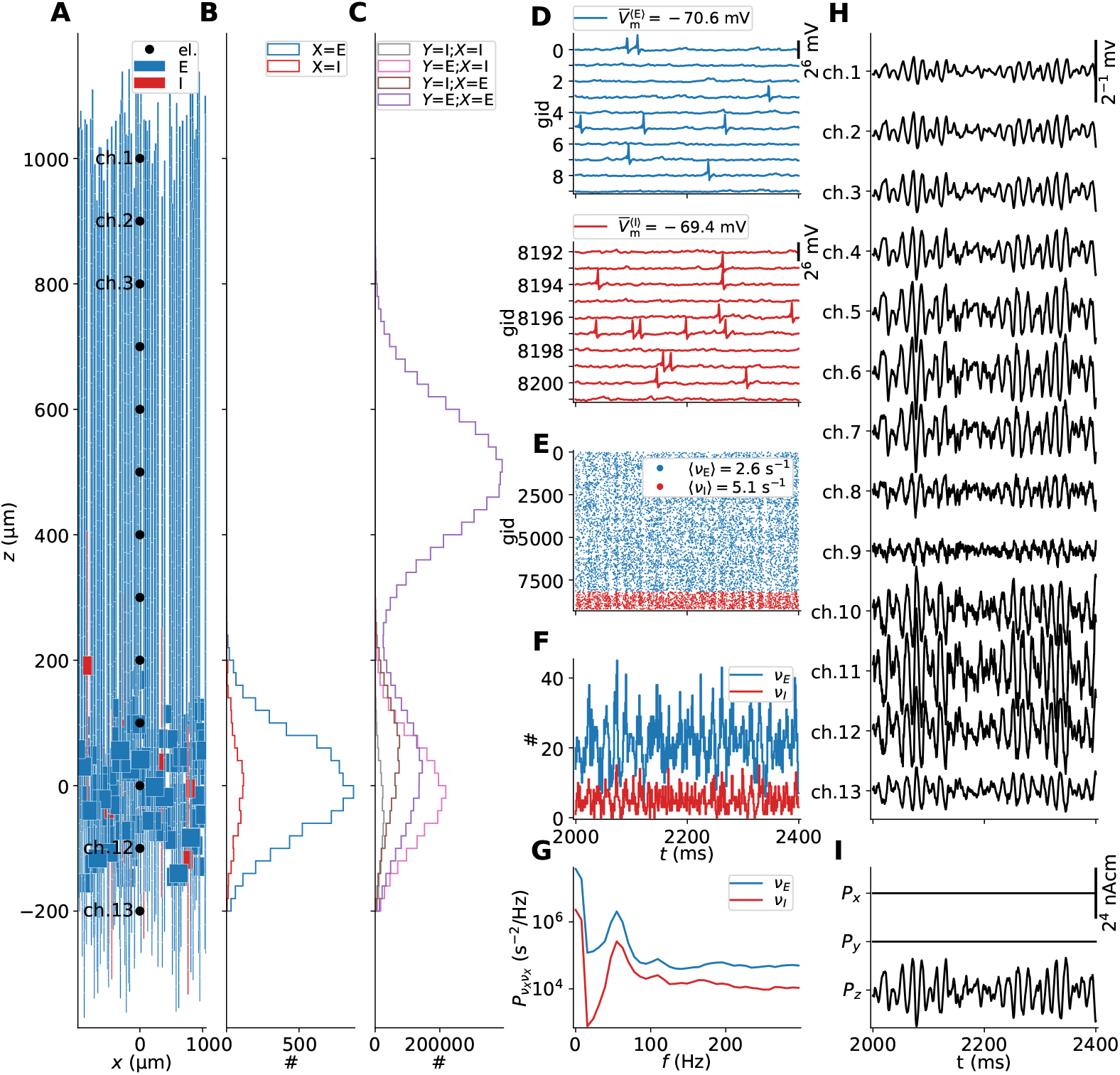
Stylized two-population MC neuron network with ground truth predictions of extracellular signals. (**A**) Neuronal populations and electrode geometry. The network is constructed of one excitatory (‘E’) and one inhibitory (‘I’) population. Only a subset of cells is shown from each population. The black point markers along the *z*-axis denote locations of electrode contact points with separation 100 µm. (**B**) Soma counts per population *X* along the vertical *z*-axis in bins of 20 µm. **(C)**Synapse counts per connection *K*_*Y X*_ along the vertical *z*-axis (bin size 20 µm). **(D)**Somatic potential traces of 10 neurons in populations ‘E’ and ‘I’. The 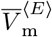 and 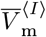 values in each legend denote median soma potentials computed from a subset of neurons in each population (*N* = 1024). (**E**) Network spike raster spanning 500 ms of spontaneous activity. The mean population-averaged firing rates are given shown in the legend. (**F**) Per-population spike-count histograms with bin size 1 ms. (**G**) Population firing-rate power spectra. (**H**) Extracellular potentials across depth (*V*_e_(**R**, *t*)). (**I**) Components of the current dipole moment (**P**(*t*)) along the *x, y, z*−axes.

### Hybrid scheme with linearized dynamics accurately captures extracellular signals of the reference network

Biophysically detailed as well as simplified networks of spiking point-neuron models can generate realistic spike train statistics of different populations. But, the presently used framework combining MC and VC models is required in order to compute meaningful extracellular population signals such as the LFP. In the hybrid scheme ([18]; Hybrid scheme for extracellular signal predictions), the simulation of spiking activity in the recurrent network(s) can be performed separately with intermediate storage of spikes, while extracellular signals can be computed via unconnected populations of MC neurons activated by synapses triggered at times as they *would have* occurred in the actual network. Using the reference recurrent MC neuron network and corresponding spike events and ground-truth extracellular signals shown in Fig 3 we can now, in contrast to our earlier study [18], test this prediction scheme in a self-consistent manner.

In this test, we record spikes trains of each neuron and ground truth extracellular potentials and current dipole moment from our reference MC neuron network to file, as well as the randomly instantiated cell locations in space and the full synaptic connectivity including synaptic placements. The resulting connectivity table includes pre- and postsynaptic neuron id, synaptic location (cell morphology coordinate and Cartesian coordinate), maximum synaptic conductance and transmission delay.

With the above information, we confirmed we can compute the intra- and extracellular signals matching the ground truth exactly, as initial conditions, neuron models and synaptic activations, etc., can be preserved in absence of actual recurrent connections (result not shown). However, one benefit of the present hybrid scheme, is that it allows simplifying the individual neuron and synaptic dynamics systematically. In particular, we shall assertain that linearized model setups can accurately capture the features of the ground truth extracellular potential (*V*_e_(**R**, *t*)) and current dipole moment (**P**(*t*)). Here, we shall account for synaptically evoked contributions to the different signals.

We first consider 4 hybrid scheme model configurations. These configurations all incorporate the same linear approximation to synaptic currents around the median somatic voltage in each reference network population as described in pt. 1 in Linear approximations to synapse and membrane dynamics. Then, we consider every possible permutation of (1) whether or not to account for changes in the effective membrane leak conductance *g*_eff_ per compartment *m* due to synaptic activity (see pt. 5 in Linear approximations to synapse and membrane dynamics), and (2) the quasi-active linearized (`biophys:lin`) and passive-frozen (`biophys:frozen`) model neuron variants representative of the ‘E’ and ‘I’ population showcased above. See pt. 2-3 in Linear approximations to synapse and membrane dynamics for details on the linearization procedure for voltage-gated ion channel descriptions.

By visual inspection of all hybrid scheme predictions in Fig 4, both model setups that account for changes in the effective membrane leak conductance (`g_eff:True`) in panels C and D accurately capture the spatiotemporal features of the ground truth signals (black lines), including signal amplitudes. The main differences seen here are that the ground truth signals contain high-frequency jitter that is not captured in hybrid scheme predictions as signal contributions by APs are not accounted for by design.

**Fig 4.**
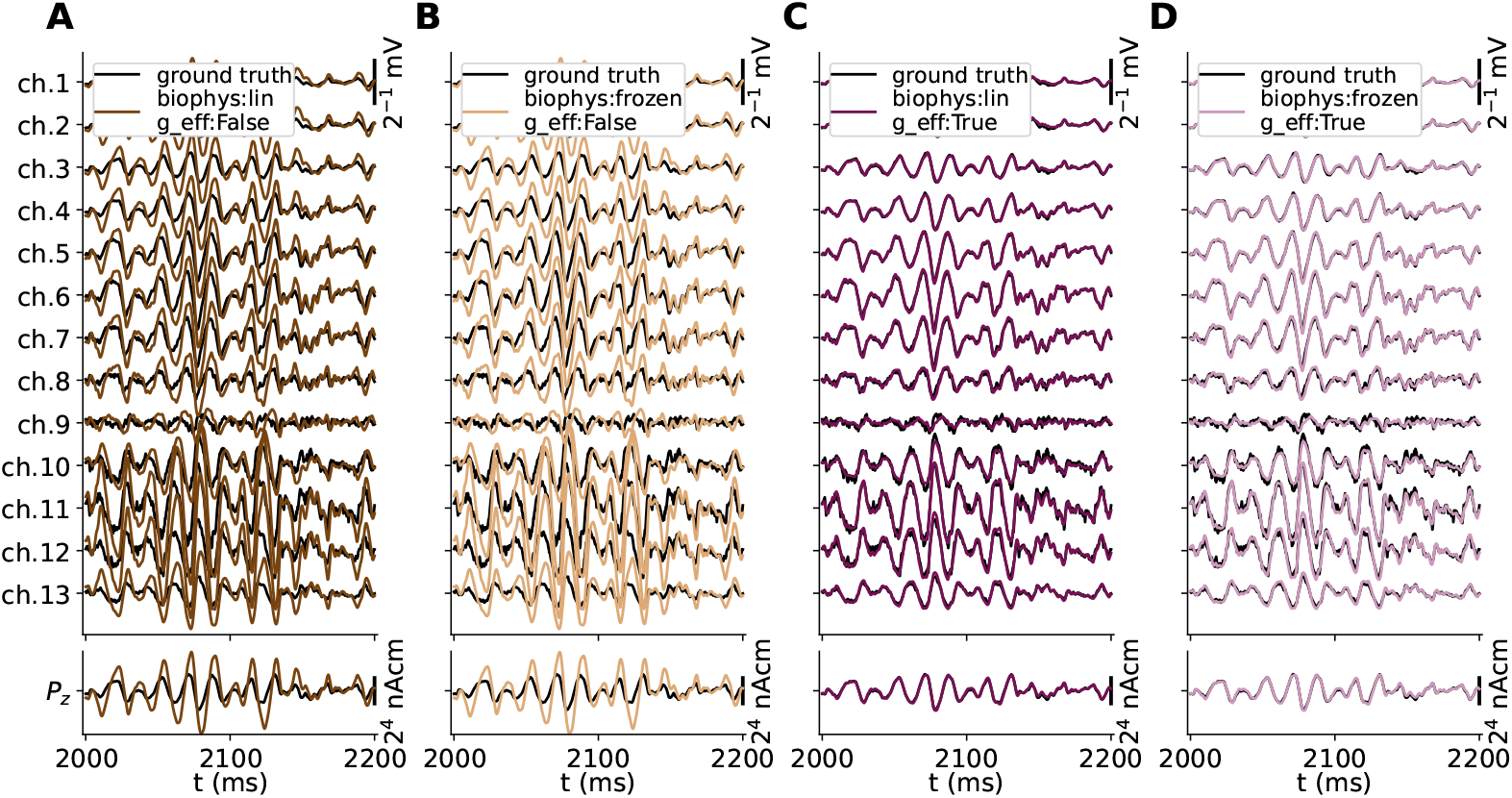
Ground truth signals vs. hybrid scheme approximations. Extracellular potential across depth (top row) and *z*-component of the current dipole moment (*P*_*z*_(*t*), bottom row) predicted from the MC neuron network model in Fig 3 (black lines) is compared to predictions made using the hybrid scheme (colored lines), using current-based synapses and neurons with either passive-frozen or quasi-active ion-channel dynamics (`biophys:frozen` vs. `biophys:lin`), ignoring or accounting for the effective membrane conductance (`g_eff:False/True` in panels A-D respectively).

Our choice of quasi-active (`biophys:lin`) or passive-frozen (`biophys:frozen`) ion-channel dynamics are seen to have remarkably little effect on the predicted signals (in contrast to somatic voltage responses). However, *not* accounting for membrane conductance contributions by synapses (`g_eff:False`), results in a clearly detrimental effect on the predicted signals (panels A and B). The most salient observation is that the approximations to extracellular potentials across depth (*V*_e_(**R**, *t*)) as well as *z*−components of the current dipole moment (*P*_*z*_(*t*)) are predicted with amplitudes that are about a factor 2 too high. The signals also appear to lag behind the ground truth signals by a few ms in the temporal domain. These effects are also observed in a preliminary report on this particular hybrid scheme model configuration [67]. A more thorough analysis and summary of the accuracies of these signal approximations are summarized below in Accurate signal predictions using hybrid scheme and deterministic kernels, and compared also to kernel-based prediction methods. For the remainder of this study, we will thus assume that methods other than the hybrid scheme must also account for changes in the leaky properties of the membrane. This is due to the effect the (effective) membrane time constant has on the integration of synaptic input currents throughout the dendrites and the resulting distributions of transmembrane currents.

### Kernels for accurate signal predictions

So far we have shown that hybrid scheme predictions incorporating linear approximations to the synapse and active ion channel currents accurately capture the extracellular potentials across depth as well as the current dipole moment. This observation implies that the relations between times of presynaptic APs and resulting spatiotemporal distribution of transmembrane currents (and therefore extracellular potentials etc.) of respective postsynaptic neurons are approximately fixed. As the postsynaptic responses can not occur before the spike times of presynaptic neurons, these relationships must also be causal. Throughout this and the next sections, we shall therefore further test the idea that extracellular signal predictions can be well represented as a linear time-invariant (LTI) causal system. Here, we shall compare filter coefficients, or ‘kernels’, obtained at the population level using two different approaches, either via the hybrid scheme setup above, or using a novel, direct, deterministic method based on the idea that the underlying distributions of cell and synapse positions, synaptic delays, linearized ion-channel, linearized synapse dynamics, and neuronal geometries provide sufficient information to estimate the corresponding causal filters. Our derivation of these deterministic kernels is described in detail in Direct kernel predictions from single MC simulations. In both cases, the kernels represent the population-averaged postsynaptic response of spike events in each presynaptic population, that is, equivalent to ‘spike-signal’ impulse response functions of the system. For corresponding signal predictions evaluated in Accurate signal predictions using hybrid scheme and deterministic kernels, the kernels are applied with population spike rates. Predictions are compared with ground truth signals generated by our reference recurrent MC neuron network (see Reference MC neuron network with extracellular signal predictions).

#### Predicted kernels using the hybrid scheme method

As discussed in [18], estimating full sets of kernels for every connected pair of pre- and postsynaptic neurons for signal predictions is intractable in large networks due to the connection count and corresponding kernel count. The study showed that averaged kernels *H*_*Y X*_ (**R**, *τ*) computed for presynaptic (*X*) and postsynaptic (*Y*) populations could accurately capture the corresponding hybrid scheme extracellular potentials by the double sum over the convolution of population firing rates and averaged kernels (see Eq 25). Here, we revisit this approach, adding also current dipole moments to the comparison.

First, we take the hybrid scheme simulation above, using current-based synapses and either variant of linearized ion-channel dynamics. We account for changes in the effective membrane leak conductance as above (`g_eff:True`). Then, ongoing spiking activity in each population is replaced by single synchronous events that allow for computing the full set of population-averaged kernels *H*_*Y X*_ (**R**, *τ*) using Eq 26. The resulting sets of kernels for predicting the extracellular potential and current dipole moment are shown in Fig 5A. Consistent with our earlier observation, only minor differences occur between kernel signals predicted using quasi-active or passive-frozen cable models. The set of kernels reveals non-trivial relationships between spikes by neurons in each population and the extracellular potential across depth due to combined effects of the cable models, synapse model, VC model, etc., and could challenge model assumptions made in other studies like space- and time-separable kernels (e.g., [20, 34, 35]) due to the effect of dendritic integration.

**Fig 5.**
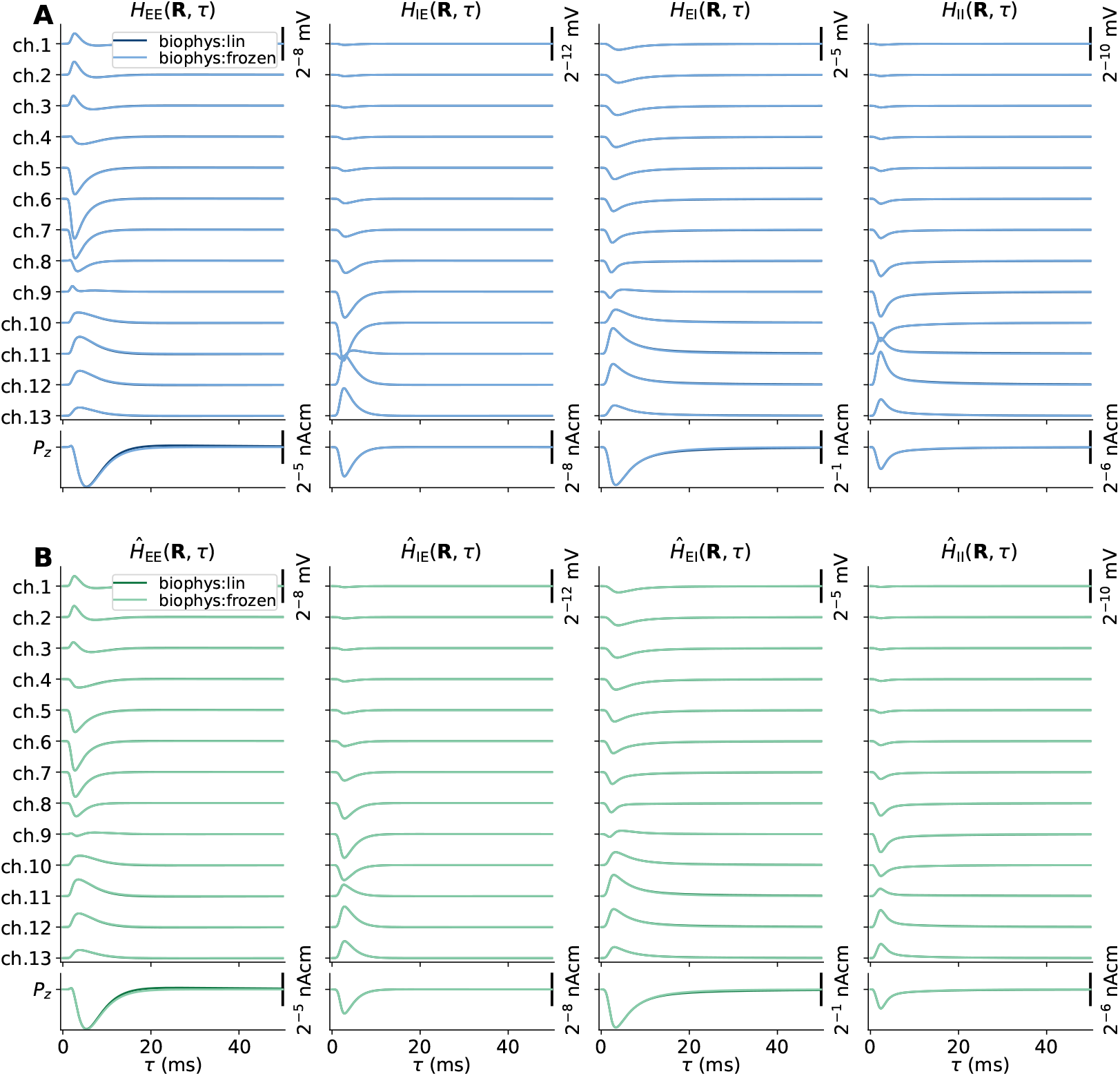
Spike-signal impulse response functions (‘kernels’) for extracellular potentials and current dipole moments. (**A**) Hybrid scheme spatiotemporal functions *H*_*Y X*_ (**R**, *τ*) for each connection between every possible pre- and postsynaptic network population *X* and *Y*, respectively. The top row kernels are computed as the spike-averaged contribution by postsynaptic neurons to the extracellular potential in electrode contact locations shown in Fig 3A, while the bottom row kernels are computed as the spike-averaged current dipole moment contribution along the vertical *z*-axis. The kernels are computed either using fully passive-frozen (`biophys:frozen`) or with quasi-active (`biophys:lin`) cable models. The kernels are truncated at time lags *τ* ∈ [0, 50 ms]. (**B**) Same as panel A, but here the kernels are computed using a computationally fast and deterministic method accounting for expectation values in terms of cell and synapse placement.

The set of kernels also allows for some insight into which connections and populations shape the extracellular signals. Here, the I to E kernels (*H*_EI_(**R**, *τ*)) have amplitudes that are ∼4-16 times those of the E to E kernels (*H*_EE_(**R**, *τ*)). Thus any spike in population ‘I’ may give a significant signal contribution from inhibitory synaptic currents in population ‘E’, even if the number of neurons in population ‘E’ is 8-fold that of population ‘I’. The dominance of inhibitory over excitatory contributions in the LFP is in agreement with previous reports (e.g., [18, 68]). It should, however, be noted that our choices of synaptic density shape functions for each pathway (*L*_*Y X*_ (*z*) defined in Table 2) may significantly affect the corresponding kernel appearances – inhomogeneous synapse densities may result in much stronger responses than homogeneous densities [41, 60, 69]. Furthermore, the direct contribution by evoked transmembrane currents on population ‘I’ can be expected to be minor, in part explained by the smaller spatial extents of the neurons and low cell count.

#### Predicted kernels using the direct and deterministic method

The kernel calculations via the hybrid scheme above rely on a number of MC neuron simulations proportional to the overall network size, and incur significant computational costs. Here we rather account for distributions and expectation values in the parameterization of the MC neuron network directly, allowing predictions of an appropriate and accurate set of kernels without instantiating network-equivalent populations of MC neurons. Described fully in Direct kernel predictions from single MC simulations, the constituents needed for these calculations are: linearized versions of the MC neurons representing each population; their distribution in space; probabilities for synaptic placements per compartment for each main connection; synaptic indegree distributions over instantiated connections for each main connection; conduction delay distribution for each main connection and the linearized synapse currents for each main connection. Typical presynaptic spike rates need to be specified as well as pairwise connection probabilities between neurons in each population. Finally, the VC model for each signal is modified to account for radially symmetric cell distributions in space (see Modified forward models for deterministic kernel predictions for details).

In contrast to the above hybrid scheme kernels shown in Fig 5A, each kernel now requires only a single MC neuron simulation to compute the population-averaged transmembrane currents following synaptic activation, and account for all other effects by a series of linear convolution operations in the spatial and temporal domains as well as a scaling by the presynaptic population size (see Direct kernel predictions from single MC simulations for details). The set of calculations results in deterministic outcomes, and are fast to compute on laptop computers while high-performance computing resources are generally required for the hybrid scheme setup. From our default parameterization of the MC neuron network (Reference multicompartment neuron networks), the resulting set of approximated kernels *Ĥ*_*Y X*_ (**R**, *τ*) for each main connections between pre- and postsynaptic populations *X* and *Y* is shown in Fig 5B. This new set of kernels appears similar to the averaged kernels computed via the hybrid scheme shown in Fig 5A, suggesting that they may be used interchangeably. The main differences appear to be somewhat reduced amplitudes of the deterministic set of kernels for extracellular potentials in panel B. Next, we shall apply our predicted kernels with corresponding population spike count histograms (‘spike rates’) for signal approximations, and compare their accuracies alongside predictions using the full hybrid scheme against the corresponding ground truth (Hybrid scheme with linearized dynamics accurately captures extracellular signals of the reference network).

### Accurate signal predictions using hybrid scheme and deterministic kernels

With the sets of hybrid scheme kernels (*H*_*Y X*_ (**R**, *τ*)) and approximated kernels (*Ĥ*_*Y X*_ (**R**, *τ*)) shown in Fig 5 panels A and B, respectively, we now convolve them with the corresponding presynaptic population firing rates *v*_*X*_ (*t*), and sum up the contributions using Eq 25. In all respects, the corresponding signal predictions shown in Fig 6 panels A-D compare very favorably with the ground truth signals generated by the reference network (Fig 3H,I). By visual inspection, neither hybrid scheme predictions (Hybrid scheme with linearized dynamics accurately captures extracellular signals of the reference network, Fig 4) nor kernel-based predictions display clearly distinguishable discrepancies from the ground truth signals in terms of spatiotemporal features and signal amplitudes, except for some high-frequency jitter associated with APs present in the ground truth data.

**Fig 6.**
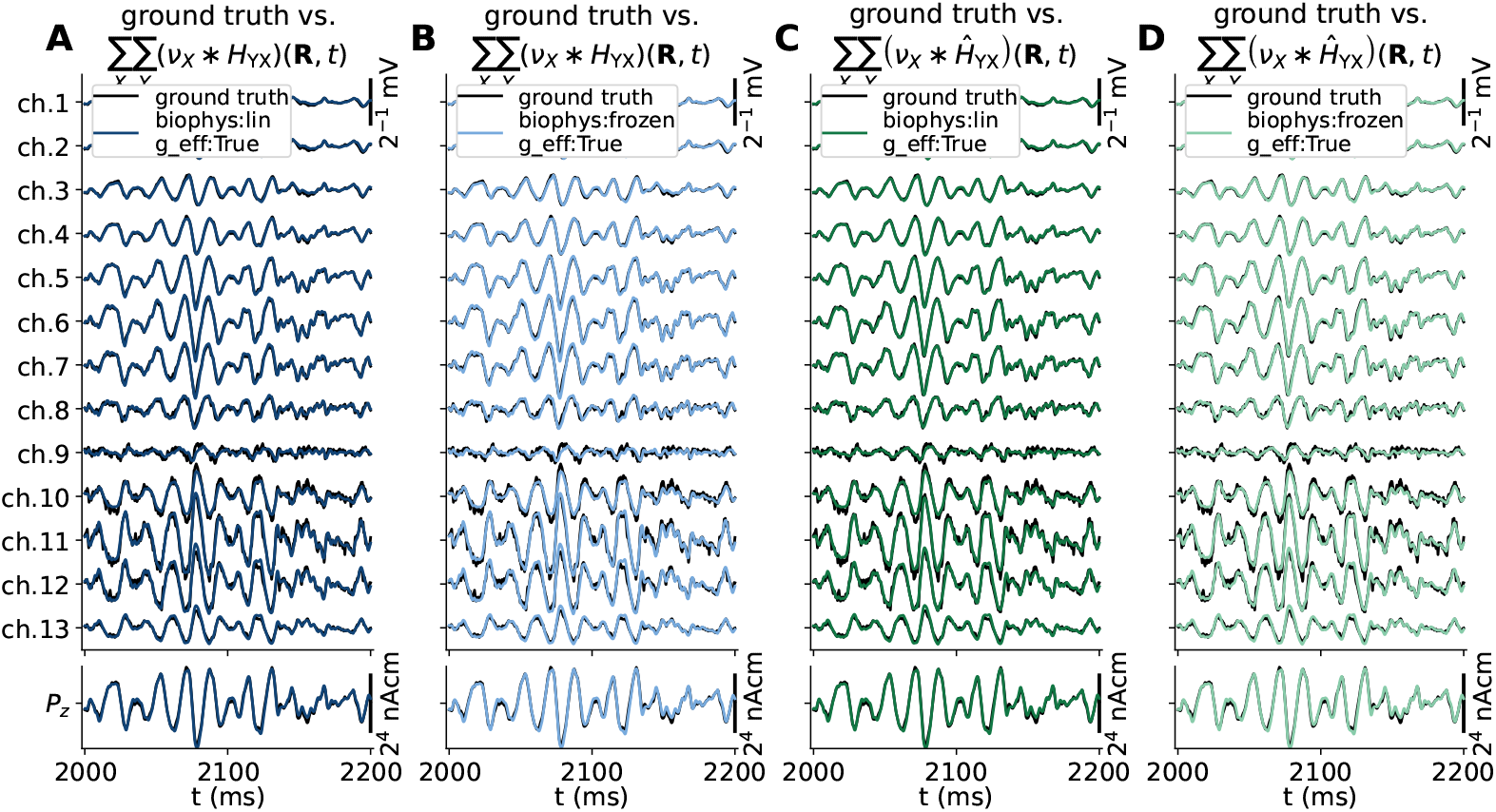
Ground truth signals vs. kernel-based approximations. (**A**,**B**) Ground truth extracellular potential (top) and current dipole moment (bottom) predicted from the MC neuron network model in Fig 3 compared to predictions made using the hybrid scheme kernels *H*_*Y X*_ (**R**, *τ*) shown in Fig 5A. The signal approximations are obtained by convolving presynaptic population firing rates (*v*_*X*_ (*t*)) with respective kernels (*H*_*Y X*_ (**R**, *τ*)) and summing the contributions. (**C**,**D**) Same as panels A and B, using deterministic kernels *Ĥ*_*Y X*_ (**R**, *τ*) shown in Fig 5B.

In order to quantify prediction accuracies, we therefore resort to comparing squared Pearson correlation coefficients (*R*^2^, Eq 38) and relative differences in their standard deviations (*r*_STD_, Eq 37) between ground truth signals and predictions. We compute these metrics not only for the ‘raw’ signals but also for low-pass (‘LP’) filtered data. Thus by attenuating the higher frequencies typically associated with presynaptic Aps present in the ground truth (see Signals and signal analysis methods for details) a somewhat improved accuracy for the different approximations can be expected. In terms of extracellular potentials and the low-pass filtered counterpart (a.k.a. the LFP), the *R*^2^ and *r*_STD_ metrics in Fig 7 confirm our visual analysis. The worst-performing configurations are hybrid scheme setups that do not account for changes in the effective membrane time constants (`g_eff:False`). All other configurations perform well in all channels except ch. 9. Nearby this depth, the sign of the signals flips due to current conservation, perhaps most evident in the dominating kernels *H*_*EI*_ (**R**, *τ*) and *Ĥ*_*EI*_ (**R**, *τ*) shown in Fig 5. Except for the ch. 9 outliers, the observed *R*^2^ and *r*_STD_ values approach 1. Fig 7 panels C and D projects median as well as the 10% and 90% percentiles of *R*^2^ and *r*_STD_ values computed across channels. Here, an overall gain in *R*^2^ is seen in all cases in the low-pass filtered data. Overall, our choice of quasi-active vs. passive-frozen membrane dynamics has only a minor effect in terms of the *r*_STD_ and *R*^2^ metrics.

**Fig 7.**
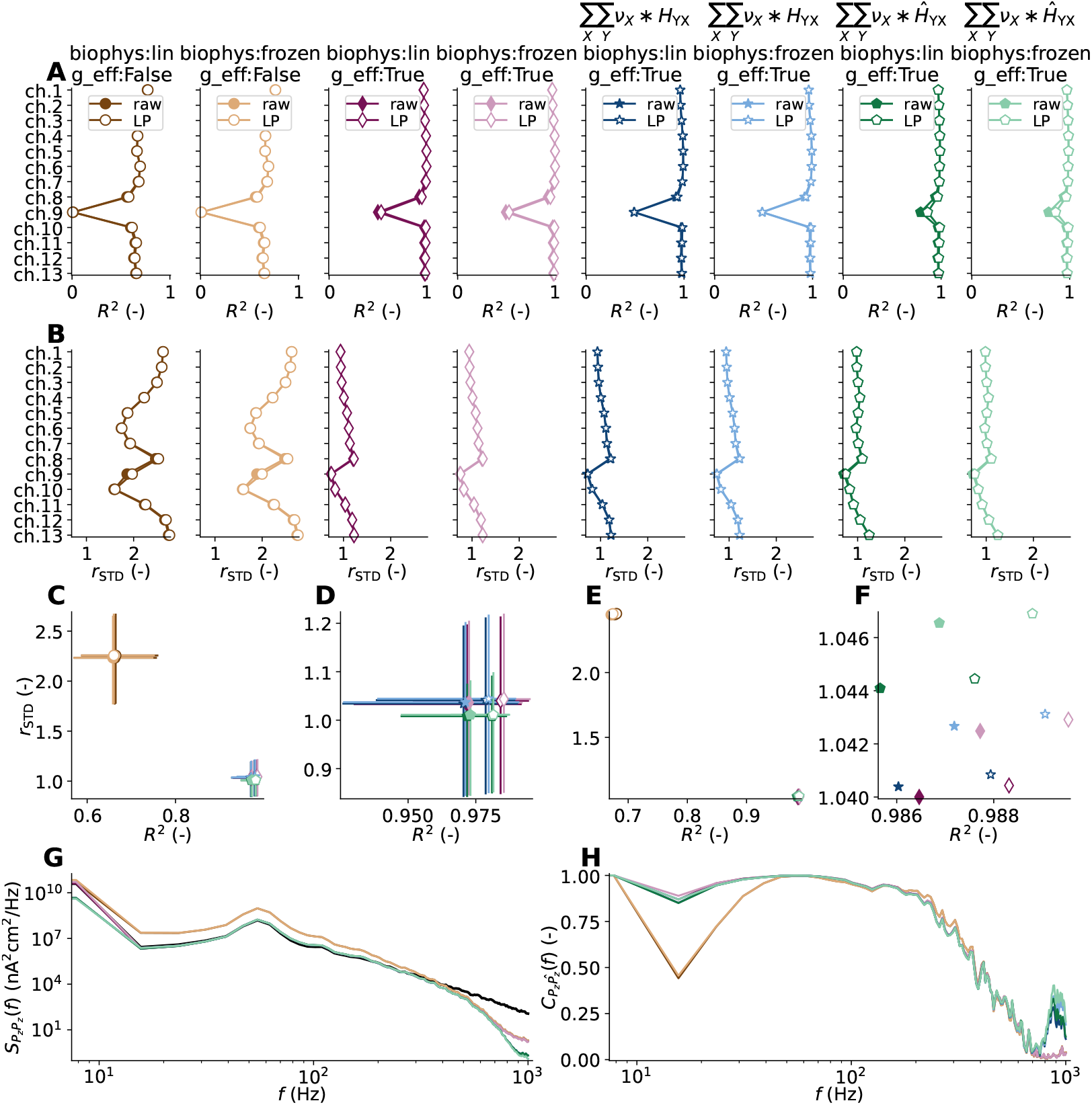
Accuracy of signal predictions vs. ground truth. For each approximation of extracellular potentials shown in Fig 4 and 6, their accuracy is evaluated in terms of the (**A**) squared Pearson correlation coefficient between approximation and ground truth (*R*^2^) and (**B**) their standard deviation normalized by ground truth signal standard deviation (*r*_STD_). The filled and white-faced markers denote metrics computed from raw and low-pass filtered data, respectively. (**C**) Aggregate *R*^2^ and *r*_STD_ values with median, 10 % and 90 % percentiles along each axis computed from extracellular potential approximations. Outliers (*<* 10 %, ≥ 90 % percentiles) not shown. (**D**) Same as panel C for predictions accounting for changes in effective membrane conductance (`g_eff: True`). (**E**) Scatter plot of *R*^2^ vs. *r*_STD_ for the different approximations to the *z*-component of the current dipole moment *P*_*z*_(*t*). (**F**) Same as panel E for predictions accounting for changes in effective membrane conductance (`g_eff:True`). (**G**) PSD of the *z*−component of the current dipole moment, comparing ground truth (black line) versus hybrid- and kernel-based signal approximations (colored lines). Same color coding as in panel A. (**H**) Coherence between ground truth *z*−component of the current dipole moment and different approximations. Same color coding as in panel A.

Our findings for the extracellular potentials are mirrored for the approximated *z*−component of the current dipole moment in panels E and F in Fig 7. All approximations taking into account the effect of the effective membrane leak conductance perform excellently, both with respect to the *R*^2^ and *r*_STD_ metrics. Similarly, the PSDs of ground truth (black curve) and different approximations to the *z*-compontent of the current dipole moment in panel G, show that the spectral signal content is well captured below approximately 300 Hz. Around similar frequencies, the corresponding coherences (cf. Eq 41) in panel H drop below approximately 50 %.

### Effect of perturbed parameters on signal predictions with deterministic kernels

Predictions of kernels *Ĥ*_*Y X*_ (**R**, *τ*) and corresponding signals rely on accurate assessments of a number of parameters. Here we choose to investigate the effect on *Ĥ*_*Y X*_ (**R**, *τ*) of mismatched time-averaged presynaptic population firing rates ⟨*v*_*X*_ (*t*) ⟩ (including that of the external population) and choice of 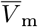 on our *R*^2^ and *r*_STD_ metrics. The unperturbed 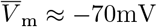 (Fig 3D), and presynaptic rates ⟨*v*_*X*_ (*t*) ⟩ are 2.6 s^−1^, 5.1 s^−1^ and 40 s^−1^ for excitatory, inhibitory and external synapses, respectively. The *R*^2^ and *r*_STD_ statistics are computed for rate-based time-series predictions against corresponding ground truth datas (Fig 3H,I). For brevity, we chose to compute these metrics only for the *z*−component of the current dipole moment (*P*_*z*_(*t*)). In our results above this term appears to be a valid indicator for corresponding metrics computed from extracellular potentials (*V*_e_(**R**, *t*)). The parameter ⟨*v*_*X*_ (*t*) ⟩ directly affects the calculation of the effective leak conductivity values *g*_eff*m*_ via Eq 29, while 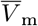 affects the linearization steps applied to voltage-gated ion channels and synaptic currents as detailed in Linear approximations to synapse and membrane dynamics. For brevity, we compute kernels employing neuron models with passive-frozen ion-channel dynamics.

The contour lines denoting *R*^2^ equal to 0.95, 0.98 and 0.99 in Fig 8A demonstrate that a relatively broad range of parameter values results in good temporal agreement between the approximated and ground truth signals. When 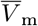 is shifted by − 10 mV the signal contributions by inhibitory synapses drop significantly as the difference to the inhibitory synapse reversal potential diminishes. If the assumed presynaptic rates are all rescaled to zero (the ratio 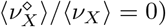, it amounts to ignoring the effective leak conductivity altogether as 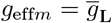. A minor gain may be seen for low-pass filtered data. The *r*_STD_ values computed across the same parameter space in Fig 8B show a more gradual dependency on each parameter. Reasonable *r*_STD_ values occur alongside the contour line labeled ‘1.0’.

**Fig 8.**
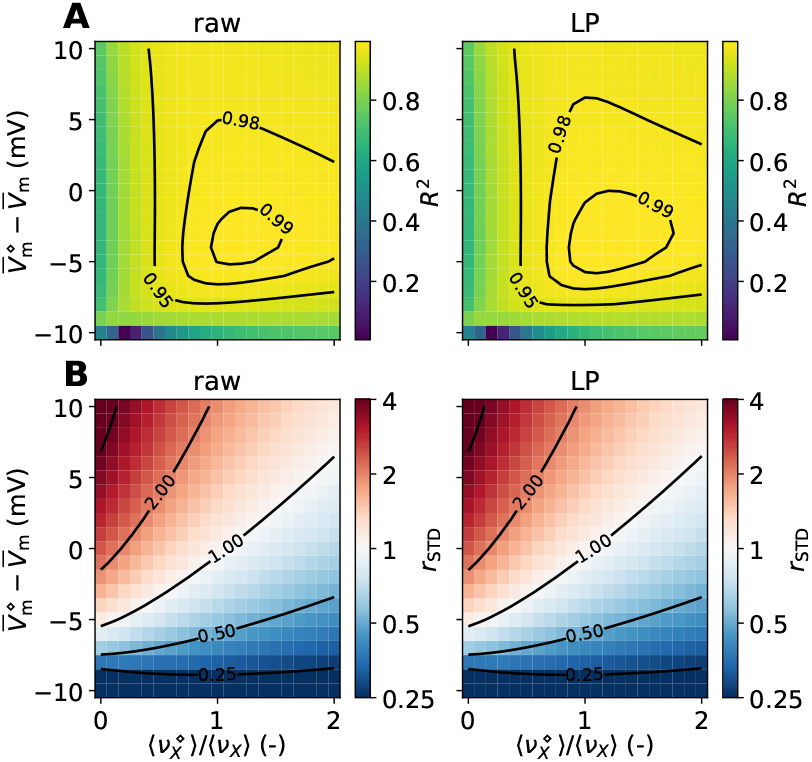
Effect of mismatched presynaptic firing rates and membrane potentials on kernel-based approximations to the current dipole moment signal. (**A**) Effect on the *R*^2^ metric computed between ground truth *z* component of the current dipole moment (*P*_*z*_(*t*)) and corresponding kernel-based approximations. For each datapoint in each panel, the kernel approximations *Ĥ*_*Y X*_ (**R**, *τ*) are computed when shifting the linearization membrane voltage by 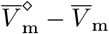 and multiplying the presynaptic firing rates by a factor 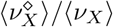. The superscript denotes perturbed values. The panels show *R*^2^ computed for kernels assuming passive-frozen (`biophys:frozen`) ion-channel dynamics. The left and right columns show *R*^2^ computed from raw and low-pass (`LP`) filtered data, respectively. (**B**) Same as panels in A but for the *r*_STD_ metric.

### Methods performance for perturbed network states

So far our Results show that fully linearized model setups can accurately approximate the ground truth extracellular signals of the reference recurrent MC neuron network. The main linearization tricks (detailed in Linear approximations to synapse and membrane dynamics) are (1) approximations of the conductance-based synapses by equivalent current-based synapses and (2) approximations of the active ion conductances by linearized versions. A crucial parameter in both cases is the choice of the postsynaptic membrane potential 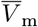 which is assumed constant. Initially, we have chosen the median somatic membrane potential averaged over neurons in each population of the reference networks. However, there are several scenarios where this assumption of near-constant postsynaptic membrane potentials can be expected to fail. This may include the presence of large-conductance synapses where synapse activation may result in significant de- and hyperpolarized postsynaptic membrane potential, as well as synchronous network states where the variance in membrane potentials may increase with the increased strength of the network-generated oscillations.

We here chose to assess the accuracy of kernel predictions and kernel-based approximations for perturbed networks in terms of modified connectivity parameters, using sets of kernels estimated directly from neuron models using passive-frozen ion-channel dynamics. For this purpose, we perturb the mean recurrent synaptic connection conductances 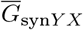 in the reference recurrent MC neuron networks by a factor governed by the parameter *J* (see Reference neuron networks with perturbed synaptic conductances for details), rerun network simulations in order to provide new ground truth data, spike trains, and somatic potentials, and recompute the set of kernels for each *J* value and derived kernel parameters. The scaling factor affects both the degree of network synchrony and overall spike rates. It also affects the kernel predictions via the updated 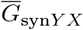 values entering Eqs. (11) and (29). *J* = 1 corresponds to our reference network model introduced above.

Our findings, summarized in Fig 9, show in panel A that increasing *J* results in increased population firing rates and increased degree of synchrony. The increased synchrony results in stronger amplitudes of extracellular potentials in ch. 2 and 11 in panel B as well as the current dipole moment in panel C. The firing rate spectra in panel D also show that the typical oscillation frequency decreases while increasing *J*, from 55 Hz to 31 Hz. Reducing *J* attenuates the firing rates and oscillations. In terms of the temporal agreement between ground-truth and approximated signals (*R*^2^, panel E), only the most synchronous activity pattern (*J* = 1.075) results in reduced performance in the upper channels. In terms of the ratio of signal standard deviations (*r*_STD_, panel F), the particular network state resulting from *J* = 1.05 yielded the worst performance. The general take home-message inferred from the aggregated *R*^2^ and *r*_STD_ values in panel G is that asynchronous irregular (AI) network states, at least for this relatively simplistic two-population network, allow for kernel-based signal predictions that well capture the corresponding ground truth signals. More synchronous activity results in reduced performance of our proposed methodology in the upper three channels.

**Fig 9.**
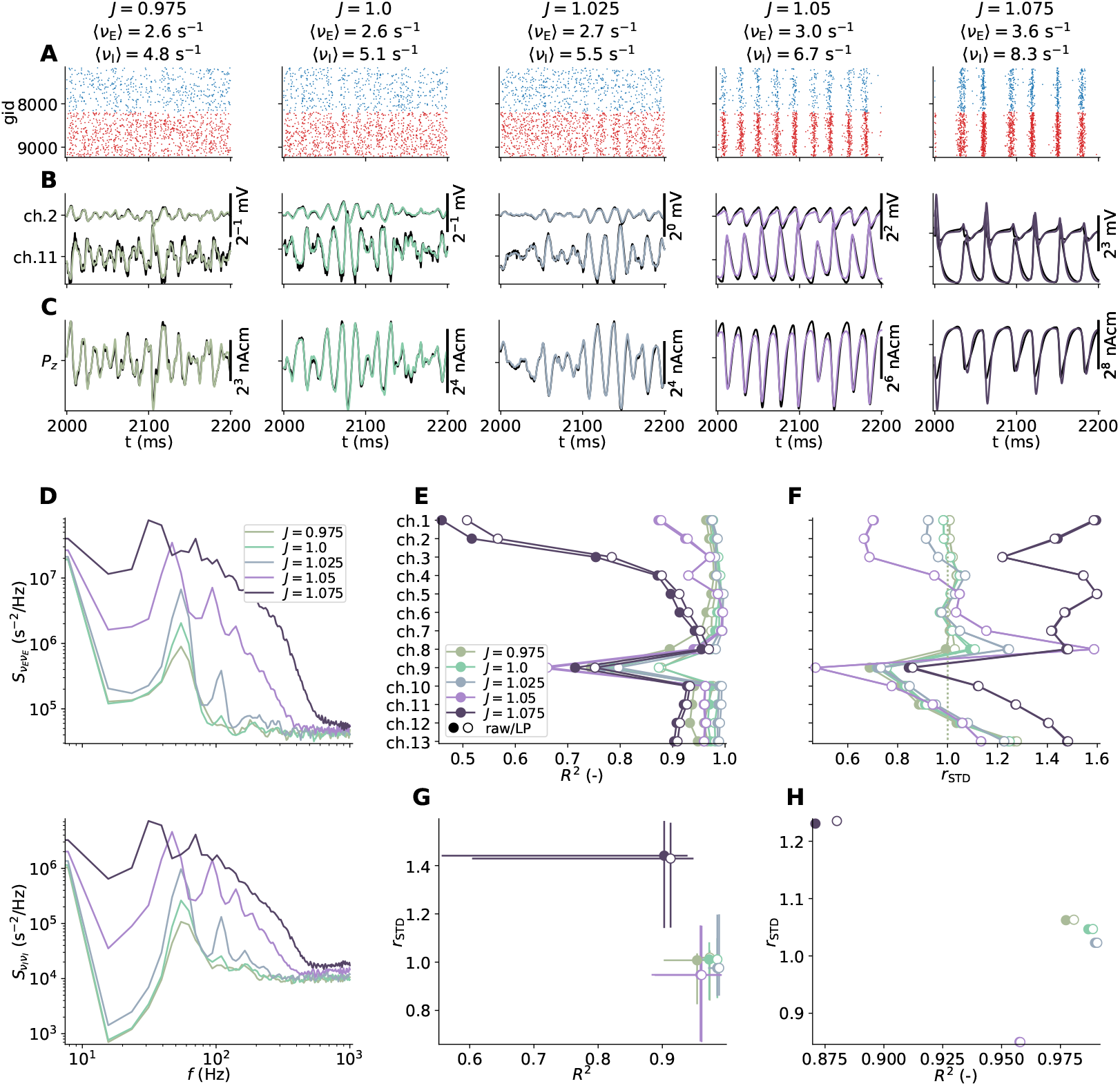
Effect of perturbed MC-network connection weights on kernel-based signal predictions. (**A**) Mean population spike rates and raster plots (*N* = 1024 spike trains in each population). The scaling factor *J* rescales all connection weights in each network simulation. *J* = 1 corresponds to our unperturbed reference network. (**B**) Ground-truth extracellular potential (black lines) and kernel-based approximations at depth of ch. 2 and 11 (colored lines). (**C**) Ground-truth and kernel-based approximation to *z*−component of current dipole moment. (**D**) Effect of rescaled connection weights on firing rate power spectra of populations ‘E’ (top) and ‘I’ (bottom). (**E**) Accuracy of kernel predictions in terms of *R*^2^ and (**F**) *r*_STD_ for kernel-based predictions of raw- and low-pass filtered extracellular potentials. Here, kernels are in each case computed using ‘biophys:frozen’ ion channel dynamics accounting for changes in the leak conductance from synaptic conductances (`g_eff:True`). (**G**) Aggregated *R*^2^ and *r*_STD_ values (median, 10 % and 90 % percentiles) across electrode channels. (**H**) *r*_STD_ vs. *R*^2^ computed for the *z*-component of the raw and low-pass filtered current dipole moment (*P*_*z*_(*t*)).

As a final test we also recompute the accuracy metrics for *P*_*z*_(*t*) in Fig 9H. Also here, reduced performances of the kernel-based method are observed for the more synchronous networks, quantified in terms of *r*_STD_ and *R*^2^. For the signals we consider, the kernel-based approach works marginally better for the low-frequency signal components, reflected in the improved *R*^2^ values over the raw signals.

### Methods performance using biophysically detailed cell models

So far in this paper, we kept the neuron and network model descriptions at a deliberately low level of complexity. However, biological neurons are commonly modeled at a much greater level of biophysical detail both in terms of geometry and in terms of the presence of heterogeneous types of ion channels, and are also used in large-scale MC neuron network simulation studies (e.g., [70]). Here we explore how well extracellular signals of neural activity can be captured using the linearization steps introduced for networks using stylized neurons, in networks incorporating biophysically detailed neuron models. For this purpose, we replace the excitatory neurons in our previous reference networks with a rat layer 5b pyramidal cell model [30], rerun network simulations to regenerate ground truth extracellular signals etc., and repeat the analyses of hybrid- and kernel-based approximations. This detailed neuron model has many more active ion channels than the ball-and-sticks neurons and may produce back-action-potential activated Ca^2+^ spikes [30]. The network parameterization is kept identical, except for an increased indegree of external excitatory input to this population in order to preserve overall firing rates (see Reference networks of biophysically detailed neuron models for details). The hybrid and kernel-based approximations rely on linearized variants of the biophysically detailed neuron model, showcased in Neuron models with linearized membrane dynamics. To emphasize on effects explained by this change of model neuron, we exclude signal contributions by transmembrane currents of inhibitory neurons in the analysis.

#### Hybrid scheme signal predictions

First, we consider the hybrid scheme setup, where spike events of the recurrent network are used for synaptic activation times in populations of neurons but without recurrent connections, and repeat the experiments first set up for the ball-and-sticks networks.

Summarized in Fig 10; If the effective leak conductance is not accounted for (`g_eff:False`, panel A,B), signal amplitudes are clearly overestimated. Predictions are in better agreement with the ground truth when the leak conductance contribution from synaptic activation is accounted for (`g_eff:True`, panels C,D). In contrast to the previous model setup, the ground truth extracellular potential signals contain prominent extracellular spike contributions, in particular across the soma-proximal ch. 9-12. In terms of choice of linearized membrane dynamics, the visual differences are minuscular. Dynamics are linearized around 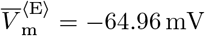.

**Fig 10.**
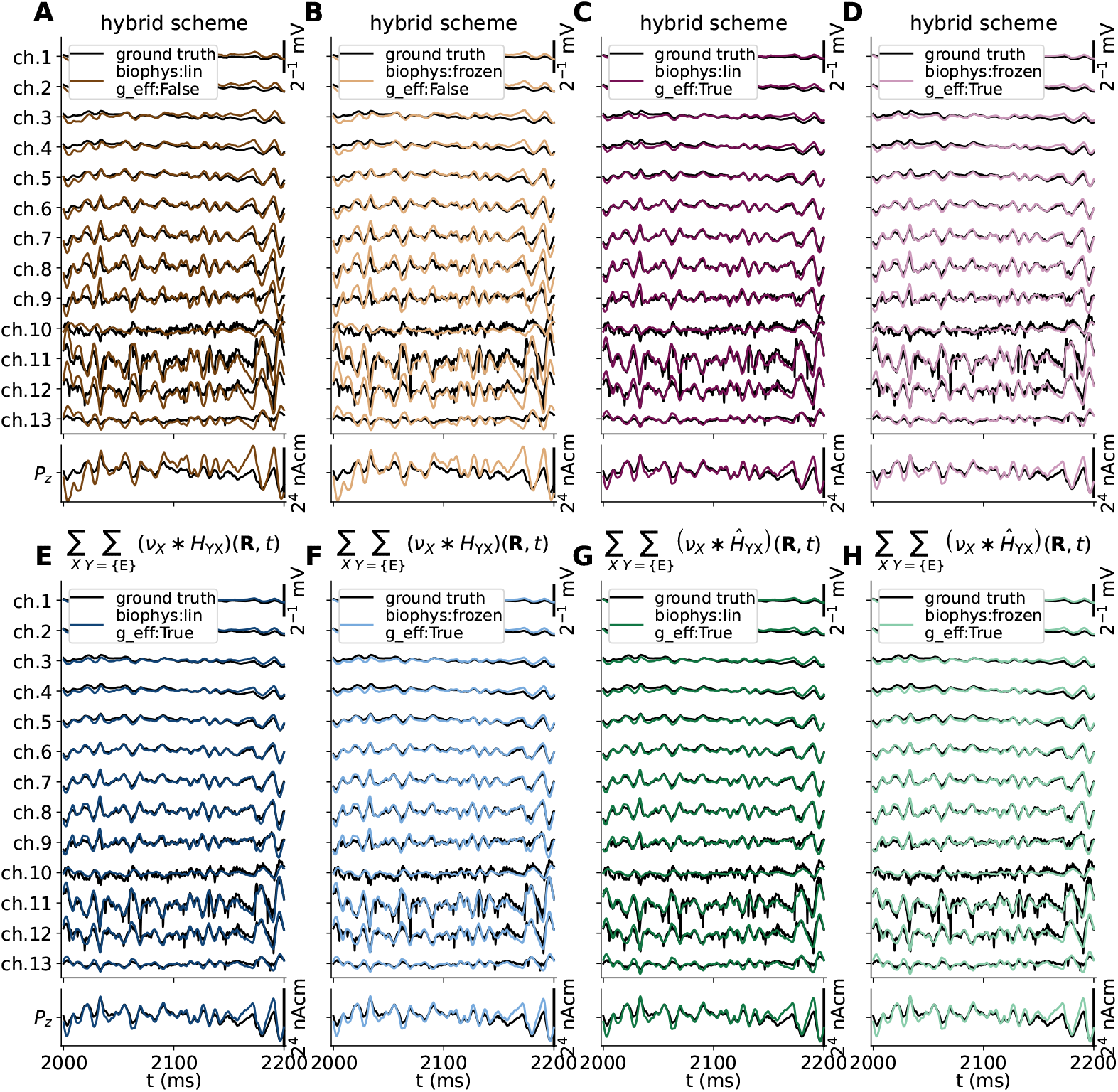
Ground truth signals vs. hybrid scheme and kernel-based approximations. (**A-D**) Same as Fig 4 and (**E-H**) Fig 6, but with the excitatory cell model being replaced by a biophysically detailed pyramidal cell model [30]. Here, only signal contributions by transmembrane currents of the updated excitatory population are accounted for.

#### Kernel based signal predictions

Next, we compare spike-to-signal impulse response functions (‘kernels’) computed via the hybrid scheme setup and the computationally fast deterministic method. The resulting set of kernels for connections onto the excitatory population in Fig 11, show that the deterministic method yields quantitatively similar kernels as the corresponding hybrid-scheme-based method. The differences can in part be explained by the fact that the hybrid implementation employs discrete synapse and cell placements in space, as they occur in the recurrent network used for ground truth signal generation, while the direct method only accounts for the underlying distributions used to set up the recurrent network in the first place. Note that with the reconstructed neuron there is a higher degree of freedom in terms of discrete synapse placements compared to the ball-and-stick neuron. The current dipole moment kernels for the E-to-E and I-to-E projections remain very similar, although visible differences now occur between the quasi-active and passive-frozen model neurons.

**Fig 11.**
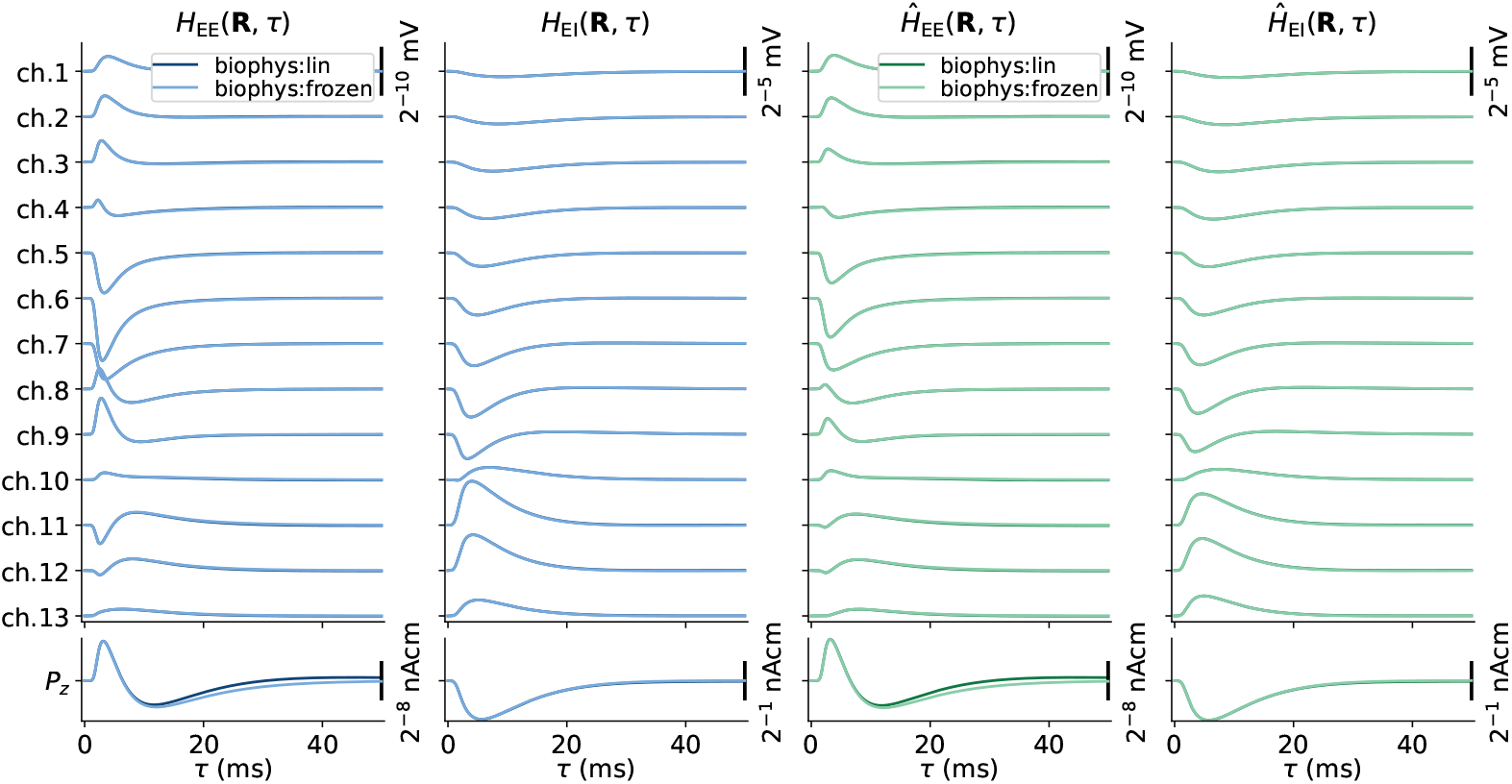
Spike-LFP impulse response function averages and predictions. Similar to panels in Fig 5, but but with the excitatory cell model being replaced by a biophysically detailed pyramidal cell model [30].

The corresponding signal predictions using these sets of kernels in combination with population firing rates are shown Fig 10 panels E-H. Similar to the hybrid scheme predictions in panels A-D, visual inspection reveals only small differences. Thus, we recompute our accuracy metrics as summarized in Fig 12 including also the hybrid scheme predictions. Similar to our initial results with ball-and-sticks neuron networks, the projected accuracies for all approximations remain clustered together if the effective membrane leak conductance is accounted for. However, the *R*^2^ metric is reduced, particularly in the uppermost channels, while *r*_STD_ is increased irrespective of signal type compared to our earlier results. The different approximations are observed to perform better in the low-frequency range as contributions by presynaptic APs in the ground truth signals are attenuated (‘LP’ vs. ‘raw’ data, respectively). The spectra and coherences comparing ground truth and approximations to *P*_*z*_(*t*) in Fig 12 panels G and H, respectively, show that signal approximations match the ground truth up to frequencies around 300 Hz.

**Fig 12.**
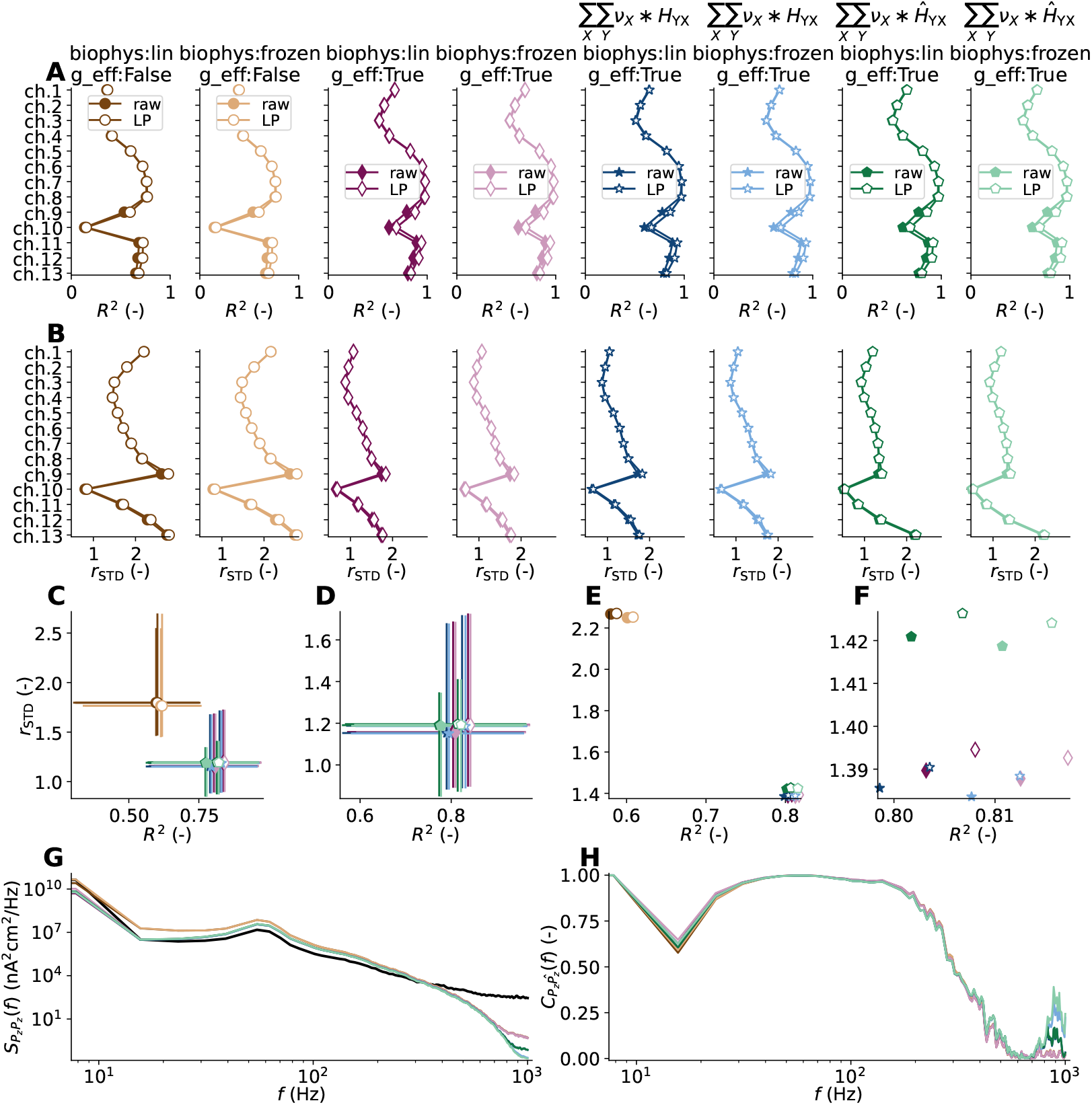
Accuracy of signal predictions vs. ground truth. Same as Fig 7, but with the excitatory cell model being replaced by a biophysically detailed pyramidal cell model [30], and accounting only for contributions by transmembrane currents of this updated excitatory population.

Overall, these observations of reduced performance compared to the ball-and-sticks cases are unsurprising, as this biophysically detailed cell model by [30] has a much more elaborate dendritic structure with many thin sections and many more degrees of freedom in terms of voltage-gated ion channels. Thus, the somatic voltage value we chose for linearized synapse and membrane dynamics may poorly represent voltage fluctuations and deviations that may be present, particularly in dendrites located remotely from the soma. Still, all approximations are able to provide excellent insight into the spatiotemporal properties of the extracellular potential and current dipole moment (and by extension EEG and MEG-like signals computed from it), more so in the low-frequency band.

For these kernel predictions we also repeated the experiment where the linearization voltage 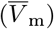 and assumed presynaptic firing rates (⟨*v*_*X*_⟩) are offset in S2 Fig. For the corresponding predictions of *P*_*z*_(*t*), a somewhat better agreement between the ground-truth and the approximated signal amplitudes can potentially be obtained by shifting 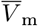 by about −5 mV.

#### Methods performance for perturbed network states

Further testing of the kernel-prediction methodology, we repeat our earlier experiment investigating the effect of perturbed conductances for recurrent synaptic connections on our proposed methodology, by introducing a variable *J* affecting 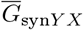 in ground-truth generating networks using the biophysically detailed layer 5 neuron model. Summarized in Fig 13, also here increasing *J* results in increased strength of network oscillations (synchrony), but the change of excitatory cell model here also results in slow synchronous oscillations with periodicity between 150–200 ms, while the oscillations in the 50 Hz range remain present. A similar emergence of slow oscillations was observed in another phenomenological network study relying on the same model neuron [71], but such activity may also arise in simplified point-neuron networks [50]. As also observed for the ball-and-sticks neuron networks, the more synchronous network states result in reduced performance of the kernel-based methodology, particularly in the uppermost channels of the extracellular potential signal (*V*_e_(**R**, *t*)). This observation may be explained by the lack of recurrent synapses in the apical tuft.

**Fig 13.**
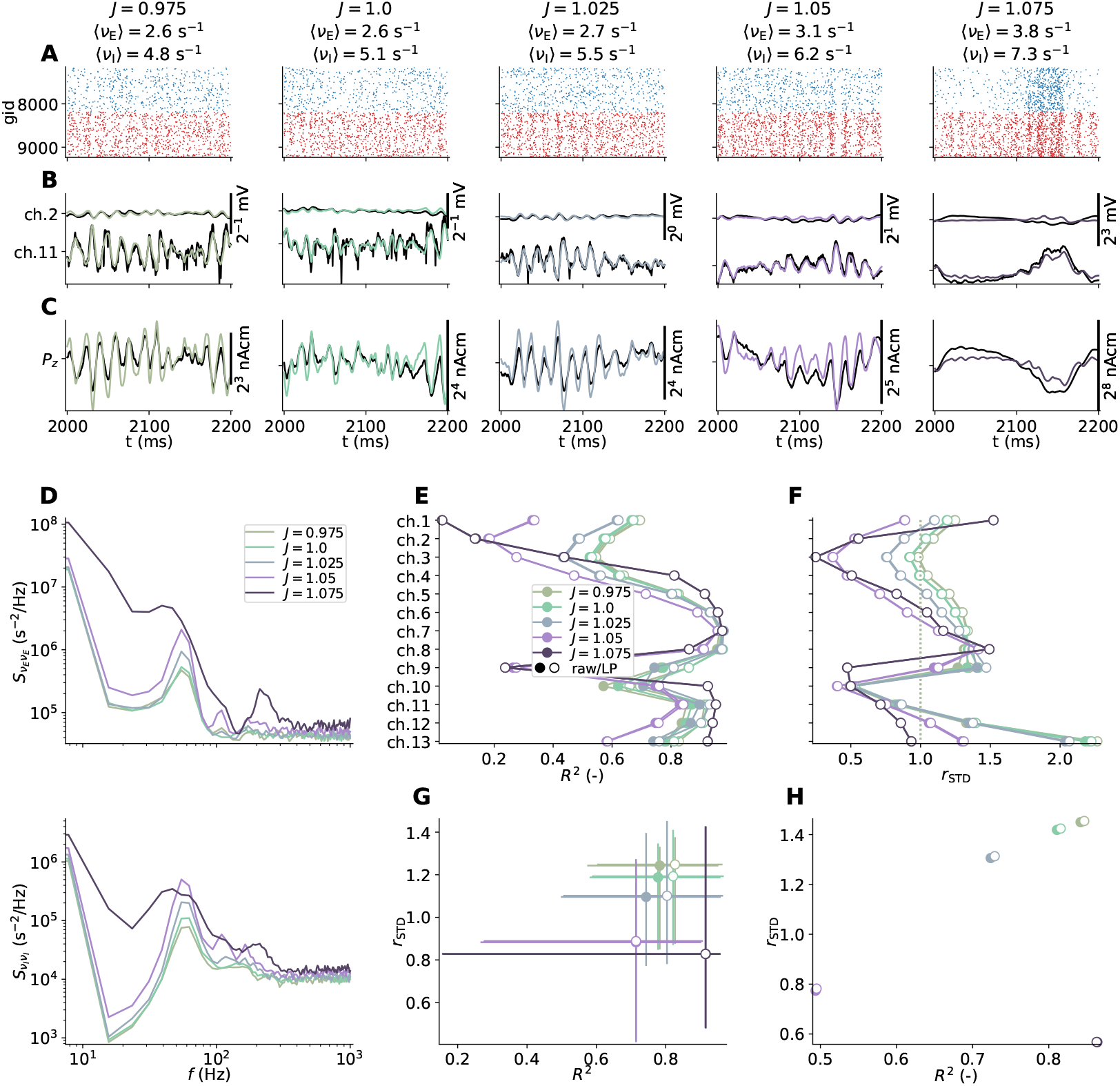
Effect of perturbed MC-network connection weights on kernel-based signal predictions. Same as Fig 9, but with the excitatory cell model being replaced by a biophysically detailed pyramidal cell model [30].

As the typical membrane voltages can be expected to vary dramatically across the elaborate geometry of the biophysically detailed pyramidal neuron, we check whether or not the accuracy of the approximated signals can be improved by varying the linearization voltage 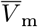 on a per-compartment basis when computing deterministic sets of kernels. For each value of *J* and corresponding network simulation, we computed the mean membrane potential per compartment across a subset of neurons (*N* = 1024) and incorporated the values when computing the set of kernels. Comparing our *R*^2^ and *r*_STD_ metrics for *V*_e_(**R**, *t*) and *P*_*z*_(*t*) for different values of *J* in S3 Fig, expose that varying 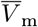 across the morphology generally increase signal amplitudes (semi-transparent markers/lines) when compared to results obtained with our earlier assumption of a constant value (opaque markers/lines). However, the overall result is inconclusive.

### Kernel-based signal predictions from point-neuron networks

Throughout Results we have demonstrated that estimates of linear spike-to-signal impulse-response functions (‘kernels’) allow for accurate approximations of different signals by convolving population firing rates with the appropriate sets of kernels and summing the contributions. So what does this allow for?

One major benefit is that spiking dynamics can with ease be modeled using recurrently connected networks employing simplified neuron representations, like leaky integrate-and-fire (LIF) point neurons and variants thereof. Recurrent network models using biophysically detailed MC neuron models (e.g., [43, 70]) are, in contrast, intrinsically more difficult to develop due to their vast number of parameters [72], are comparably slow to simulate even on large-scale high-performance computing facilities, less amenable for analytical analysis, and henceforth difficult to constrain into reasonable network states resembling experimental data. Point-neuron networks mediate all of these important issues. In addition, systematic reductionist approaches applied to MC neuron networks allow for capturing their spiking dynamics in equivalent few-compartment or point-neuron networks [73–75]. But point-neuron networks do not allow for computing the distribution of transmembrane currents in space needed for signal predictions, as all in- and out-going currents sum to zero in a point [18]. Using our direct and deterministic method we can, however, predict sets of kernels *Ĥ*_*Y X*_ (**R**, *τ*) for each connectivity pathway via single MC neuron simulations in order to compute extracellular signals from simplified networks. While reduced networks may not predict identical spike trains as the corresponding fully detailed networks, their main statistics (rates, spectra, correlations, etc.) should be preserved, implying that kernel-based signal predictions from rates remain applicable.

As a proof of principle of this methodology, we constructed a point-neuron network of the same size as our reference MC-neuron networks and fit its parameters in order to mimic our reference network’s averaged firing rates and rate power spectra shown in Fig 3 (see Leaky integrate-and-fire (LIF) point-neuron network for details). Showcased in Fig 14, the point-neuron network state is asynchronous and irregular (AI) with some oscillations present in the corresponding spike count histogram (panel A), similar to our reference network. We here also showcase the different signal contributions by each pathway (E-to-E and so forth) in panels B-E, using the set of kernels displayed in Fig 5B and discussed in Predicted kernels using the direct and deterministic method. The summed contributions are shown in panel F. Here, there are no ground truth signals to compare to directly, but the extracellular potential signal varies across time and space in an expected manner, and closely resembles the signals obtained by the I-to-E pathway. Signal amplitudes are also in the expected ranges set by our MC-neuron network simulations.

**Fig 14.**
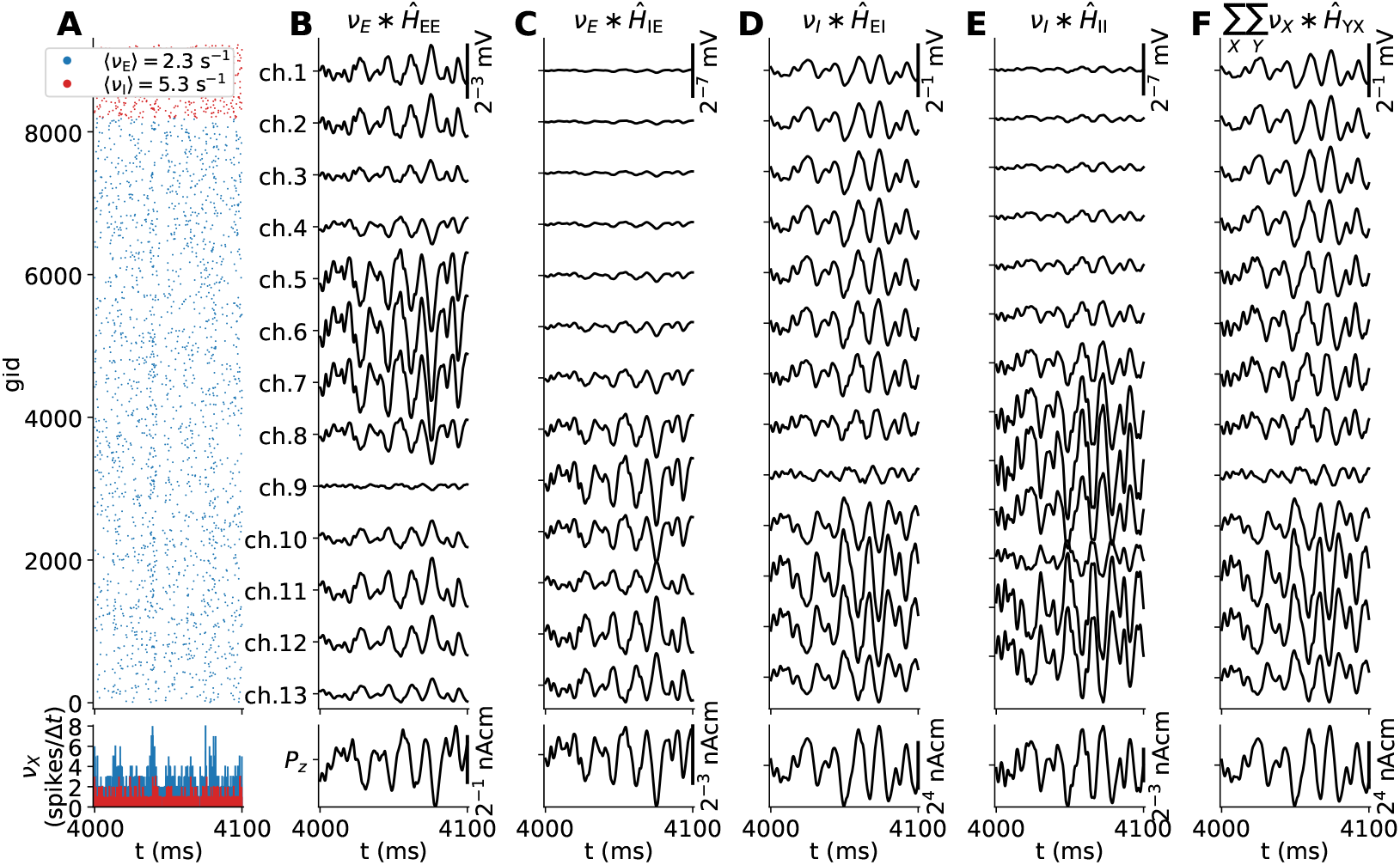
LIF network spiking activity and forward-model predictions. (**A**) Spiking activity and average spike rates of the excitatory (E) and inhibitory (I) populations of a point neuron network simulation (top), with spike counts in bins of width Δ*t* (bottom). (**B-E**) Contributions to the extracellular potential (top) and current dipole moment (bottom) by the E to E connection, E to I connection, I to E connection, and I to I connection, respectively. The signals are equivalently computed as the convolution between the presynaptic population spike count histogram and corresponding signal kernel approximations using a FIR filter implementation concurrently with the spiking simulation. The kernels used are shown in Fig 5B. (**F**) Sum over signal contributions in panels B-E.

As a final remark, we here compute the firing rates and signals ‘live’ while the network simulation is running. To reiterate, the kernels are always causal, that is, equal to zero for any time less than the minimum conduction delay in the network, and of finite duration. This causal relationship allows for treating the sets of kernels as finite-impulse-response (FIR) filter coefficients, which are here applied via a custom network node that receives incoming spike events from each population while the simulation is running and outputs continuous signals representing the temporally filtered spike events. For this purpose a FIR filter network node is implemented for the NEST simulator [52, 53] via the NESTML description language [62, 63] as detailed in NESTML FIR filter extension. This network node is also reusable for other spiking networks in NEST.

## Discussion

### Summary of findings

The main results presented throughout this paper can be summarized as follows: First, an assessment of the validity and limitations of different prediction schemes for extracellular signals from biological neuronal network models assuming linearity between times of presynaptic action potentials (‘spikes’) and corresponding extracellular signals. The signals mainly occur due to evoked transmembrane currents on the postsynaptic neuronal populations. Our finding is that the linearity assumption is valid if all contributions from the linearized membrane and synapse conductances are accounted for, resulting in accurate signal predictions.

Secondly, identification of the critical role of the effective membrane time constant due to persistent activation of recurrent and external synapses on predicted signals. We found that simply approximating conductance-based synapses by current-based synapses without accounting for the time-averaged synaptic conductances resulted in overestimated amplitudes and poorer temporal accuracy of the approximated signals.

Third, a new, fast and accurate method to compute averaged spatiotemporal spike-to-signal impulse response functions (‘kernels’) for connections between pre- and postsynaptic populations, by accounting for distributions of cells and synapses in space, linearized synapse, and membrane dynamics, overall connection probabilities, distributions of synapses per instantiated connection, and connection delay distributions. As the sets of computed kernels are causal and linearly map population spike events to the corresponding signals, it allows for efficient signal predictions as in a linear time-invariant (LTI) causal system, that is, by treating the sets of spatiotemporal kernels as finite impulse response (FIR) filter coefficients applied to corresponding firing rates of presynaptic populations. The kernel-based predictions are as accurate as a hybrid scheme explicitly accounting for neuron and synapse placements in space [18], but significantly faster. The proposed methodology accounts mainly for signal contributions resulting from synaptic activations, explaining a large fraction of the low-frequency components of extracellular signals (≲ 300 Hz).

We developed and evaluated the methodology based on recurrently connected reference networks of MC neurons. For simplicity, we initially opted for phenomenological ball-and-sticks MC neuron models with active voltage-gated ion channels distributed all over, representing each population of excitatory (E) and inhibitory (I) neurons. Synapses are conductance-based. We show that the proposed methodology is feasible with perturbed network states, as well as for cases where populations are replaced by biophysically detailed neurons [30] at a level of detail similar to neuron models implemented in high-profile biophysically detailed network modeling efforts (e.g., [43, 70]).

As a final proof of principle for the kernel-prediction methodology, we apply a suitable set of kernels for forward-model predictions from spiking activity in a spiking point-neuron network model. For this network model, the kernels are applied via a FIR filter network node receiving presynaptic spike events applying the filter coefficients for continuous signal predictions during the course of the simulation. The resulting signals resemble corresponding ground truth signals of the reference MC neuron network.

### Kernels versus other estimation methods

The sets of spike-signal kernels we compute using our proposed methodology should not be confused with corresponding spike-triggered averaged signals (e.g., [76, 77]), which are intrinsically affected by ongoing network activity, that is, spike train correlations, as previously shown in [18]. Even if both are linear measures, the spike-triggered averaged signal will most likely be non-causal and depend on the network state, unless the spiking activity of the trigger neuron is approximately uncorrelated with the ongoing activity. The latter scenario may occur for instance for spontaneous activations of neurons in one brain region (e.g., thalamus) projecting to another area (e.g. somatosensory or visual cortex, see [78–80]). This so-called monosynaptic, also referred to as unitary (e.g., by [81]) extracellular response is recently modeled in detail [82], then using conductance-based synapses but with passive membrane time constants fitted to available experimental and published data. A similar effort to compute such responses in the hippocampus was recently published [83]. Fitting such responses to spatiotemporal kernel shape functions for excitatory and inhibitory presynaptic units in order to compute LFP signals in point-neuron network models has been proposed [19]. The sets of kernels we compute do not assume a particular shape, but are derived from the biophysics of the neurons and network, and can be recomputed for other networks and populations.

Other, even simpler estimation methods for extracellular potential time series was proposed by [34], recently extended to EEG signals by [84], by approximating signals by weighted and time-shifted sums of excitatory and inhibitory synaptic currents measured in the network simulation. In contrast to the hybrid- and kernel-based approaches considered here, these simplified approximations do not explicitly account for any effects on the predicted signals from the neuronal morphologies, ion-specific channels and the VC forward model. They also do not account for any signal variation in space except if combined with some position-dependent scaling factor, and the physical units of the predicted signals can be considered arbitrary. These simplified schemes may still be considered a major improvement over *ad hoc* approaches equating firing rates or averaged somatic potentials to extracellular signals [34, 84]. In the case of scalp EEG and MEG signal predictions mainly the current dipole moment components normal to the cortical tissue surface may be predicted with reasonable accuracy and be combined with an appropriate head forward model [60], allowing for respective signal predictions along the scalp’s surface.

With the recent advances in the machine-learning (ML) field such as deep learning [85], a fair assumption is that also ML methods can infer linear/non-linear relationships between e.g., network spikes and extracellular signals if subjected to enough observations for training the algorithms. Input-output dynamics of neurons can be captured by different deep artificial neural network (ANN) architectures [86, 87], and one could likely extend such models for extracellular signal predictions. One recent study proposed deep convolutional neural networks for approximated EEG signal predictions from spike rates [84]. Linear filter-based models have also been proposed for LFP signals [18, 19, 21]. In contrast to our proposed methodology where kernels mapping population spike rates to extracellular signals are inferred from the biophysical description and parameters of the biological neuronal network itself, deep learning and related algorithms generally require experimental or model data for training. In the present context, the avenue of using ML-based methods to predict kernels from biophysical network parameters is obfuscated. Using ANNs, it was recently shown that model LFP signals contain information about underlying network parameters [38]. For mechanistic models of biological neurons and circuits, one main issue is determining suitable parameters for viable model output. Here ML-based methods such as deep neural density estimators may be used for investigating such vast model parameter landscapes [88].

### Extensions and future works

One main novelty reported here is the proposed method for directly computing kernels that facilitate efficient calculations of extracellular signals from population spike rates, as well as a reference implementation in the Python package LFPykernels. We applied this framework to quite simplified two-population recurrent networks. The framework is, however, applicable to networks with many more populations. One could for instance mimic the laminar topology of cortical microcircuits, where each layer consists of different populations representing the heterogeneous types of cells within each layer as in [70] and [43]. Based on available anatomical and electrophysiological constraints either from experiments or detailed models themselves, signal kernels of interest can then be computed for the different connections independently of simulations of recurrent network spiking activity. The latter step may then even use simplified neurons (e.g., spiking point neurons) thus negating the need for high-performance computing facilities. Extracellular signal predictions can be incorporated in the running simulation as we have demonstrated here (see Kernel-based signal predictions from point-neuron networks), or after simulation by computing population spike rates from recorded spike events, filter these and sum up all contributions.

While we have here mainly focused on the methodology and less on overall simulation speeds, we note that potential speedup can be of several orders of magnitude. The typical simulation times for the recurrent network with biophysically detailed pyramidal neuron models (see Methods performance using biophysically detailed cell models) we observe are around 4400 s multiplied by 1024 physical cores for 12 s of biological time on the high-performance computing resource, while the corresponding set of kernel predictions take around 150 s on a laptop computer using a single physical core (see Data availability and replicability for details). This number can potentially be reduced substantially if the numerical integration of Eq 35 on a per-compartment basis can be replaced by a closed-form (the MC neuron simulations of transmembrane currents are quite brief). Further reductions in prediction times may involve other trivial parallelization schemes, as kernels for different connections can be computed fully independent of each other, which is also the case for different spatial components of each spatiotemporal kernel. Code acceleration using for instance Numba^2^ or Cython^3^ may also help in this respect. Simulation times and resources required for spike times in equivalent networks of the same size using simplified neurons (i.e., few-compartment and point-neuron models) are also substantially less compared to the biophysically detailed case. For the point-neuron network incorporating the FIR filter operations used here, the respective network build and simulation times were around 8 s and 235 s with single-threaded execution on a laptop. Thus the serial time to solution is reduced by a factor ∼10^4^ compared to the MC network simulation. Hence, the avenue of biophysics-based forward model predictions of extracellular signals in large-scale networks with millions of spiking point neurons and beyond (e.g., [44]) is opened.

In its present form, there are multiple scenarios where our proposed kernel-prediction methodology could use either further development or validation. Presently we investigate the method for networks with a columnar (cylindrical) organization and no distance-dependency for connections in terms of connection probabilities, synaptic conductances, and axonal transmission delays within the column-like geometry. Large scale recurrent network models with (lateral) distance-dependent connectivity and periodic boundary conditions spanning multiple mm of cortical area has been proposed at various levels of description (e.g., [36, 43]), but so far our proposed kernel prediction method is neither developed for nor validated against such models. So far, such lateral distance-dependent connectivity was accounted for in a phenomenological kernel-based prediction model [37], and for an experimentally derived kernel-based method [19].

Furthermore, we assume recurrent networks with static connection weights. But synapses may be subject to various weight dynamics such as short-term plasticity (STP) with activity-dependent facilitation and depression, stochasticity, spike-timing-dependent plasticity (STDP) (see e.g., [89]), as well as structural plasticity [90]. Out of these, stochasticity is perhaps easier dealt with if probabilities of synaptic activations are known and independent of activation rate by scaling the corresponding kernel amplitudes accordingly. Weight changes due to STP are mainly governed by presynaptic activation intervals hence the average connection weights for kernel predictions can be determined for known averaged presynaptic rates. STDP may be harder to account for, but due to the much longer time scales for weight updates, the option to monitor connection weights during the course of simulation could allow for recomputing kernels and applying them to each simulation segment.

In terms of signal predictions in network models incorporating recurrent connections with external populations (representing other areas or nuclei) or interactions with the external world (e.g., mimicking closed-loop experiments), the present framework for direct kernel predictions could well account for the additional signal contributions. Exemplified by a putative network model of the thalamus and somatosensory cortex, representative sets of kernels must initially be computed for presynaptic spike events of thalamocortical projection neurons targeting subsets of cortical populations given knowledge of the corresponding anatomy and biophysics. Applying the additional sets of kernels with presynaptic spike events or spike rates for signal predictions would then account for locally evoked signal contributions by remote activity, without affecting network activity itself. In case synaptic weight updates (via STDP for instance) takes place, the kernels may require recalculations as suggested above.

Our analyses also demonstrate that the accuracies of kernel-based signal predictions versus corresponding reference signals can be expected to drop when the degree of synchrony and/or firing rates in the network increases (see Methods performance for perturbed network states), which we observed by rescaling recurrent synaptic conductances. Our reference network generates various-strength oscillations in the gamma range (∼ 55Hz) when driven by external fixed-rate Poisson processes, and we obtained also slow synchronous oscillations in case of the biophysically detailed neuron network. We expect to observe similar detrimental effects on prediction errors for networks with non-stationary activity. Such non-stationarities may include up-and-down states [91] or result from variable-rate external drive (e.g., representing sensory input), as the choice for membrane potential when linearizing synapse and ion-channel dynamics may indeed affect the kernel predictions. Our results of setting the linearization voltage value on a per-compartment basis are inconclusive, however, but this idea should be explored further in the future. Still, our hope is that the kernel-prediction methodology can still give excellent qualitative insight into extracellular signals from networks expressing non-stationary behavior.

Contrary to our starting point, recurrent MC neuron networks, forward model predictions from recurrent point-neuron networks pose a potential challenge due to their inherent lack of detail. Their descriptions may contain no spatial information even if the network is supposed to mimic a particular brain area, such as the generic somatosensory cortex column model proposed by [49], representing the local circuitry under a 1 mm^2^ patch of the cortical surface. To compute extracellular potentials from this model spatial information in terms of neuron geometries and depth-dependencies for synaptic placements should be determined based on available anatomical data (see [18] for details). Similarly, the present kernel predictions require MC neuron models representative of each population, and statistical distributions describing placements of cells within each population in space, placements of synapses across the neuron models for each pre and postsynaptic population, numbers of synapses per instantiated connection. Other parameters may (or may not) be derived from the point-neuron network description, such as conduction delay distributions and synaptic parameters. Some may be derived from its activity, such as population firing rates. As such, multiple concurrent efforts aim to amass such anatomical and electrophysiological detail for different brain regions and species with corresponding tools for enquiring the data (see e.g., [92–96]). Such data may be used to derive suitable kernels.

For rate-based frameworks aiming to explain activity in terms of population firing rates in finite-sized populations (see e.g., [45–48]), special attention should also be taken. Unless the rate-based models are derived using bottom-up approaches, in contrast to heuristics or inferred statistically, for instance via dynamical causal model frameworks [97], use of our proposed kernel prediction scheme also necessitates specifying parameters such as population cell counts and pairwise connection probabilities. Otherwise, the resulting kernel amplitudes can be considered arbitrary. If such parameters indeed can be determined, we do not see any principled reasons why one could not apply the kernels with continuous population rate predictions as we already demonstrated with temporally binned population spike rates computed from spiking networks.

Further extensions of our kernel estimates for continuous neural fields equations aiming to explain activity across space [98] should be based on and validated via the aforementioned laminar network models incorporating lateral distance-dependent connectivity routines. For discretized spiking point-neuron network models with distance-dependent connectivity, [99] derived corresponding neural field equations. Developments in this direction are required for simulation frameworks such as ‘The Virtual Brain’ (TVB [100]) aiming to relate firing rates across brain areas also with extracellularly recorded signals such as the EEG, as well as similar tools aimed towards clinical use [101].

Finally, we have considered only postsynaptic contributions from synaptic activations to signals predicted using the hybrid scheme or kernel-based methods. These approaches are therefore better able to capture the low-frequency parts of the signals as most clearly demonstrated in our simulations using the biophysically detailed layer 5 pyramidal neuron which resulted in clearly visible extracellular spikes in the ground-truth extracellular potentials. One could potentially account for signal contributions by presynaptic events such as somatic APs, backpropagating APs, Ca^2+^ and NMDA spikes by computing and superimposing the extracellular signatures of each event to the signals considered here, in case the network model accounts for times of such events. Taking such steps would result in non-causal kernel contributions and would require additional validation against network models using biophysically detailed neuron models expressing such phenomena. It should however be feasible to incorporate and could improve the accuracy of the present implementation around frequencies where spike contributions may dominate in the extracellular signals.

## Conclusion

Many of the research successes in the physical sciences have come from an interplay between modeling and experiments where predictions between physics-based candidate models have been systematically compared with experiments in an iterative back-and-forth loop. This approach is sometimes referred to as the ‘virtuous loop’ or circle [102]. For large-scale network models in the brain, this approach has until now been hampered by the lack of physics-based forward models able to predict mesoscopic and macroscopic brain signals like LFPs and EEGs [6]. We believe that the kernel-based approach presented here could be an important step forward for making such model-based predictions feasible, thus paving the way for use of the virtuous loop also in large-scale network neuroscience.

## Supporting information

**S1 Fig.**
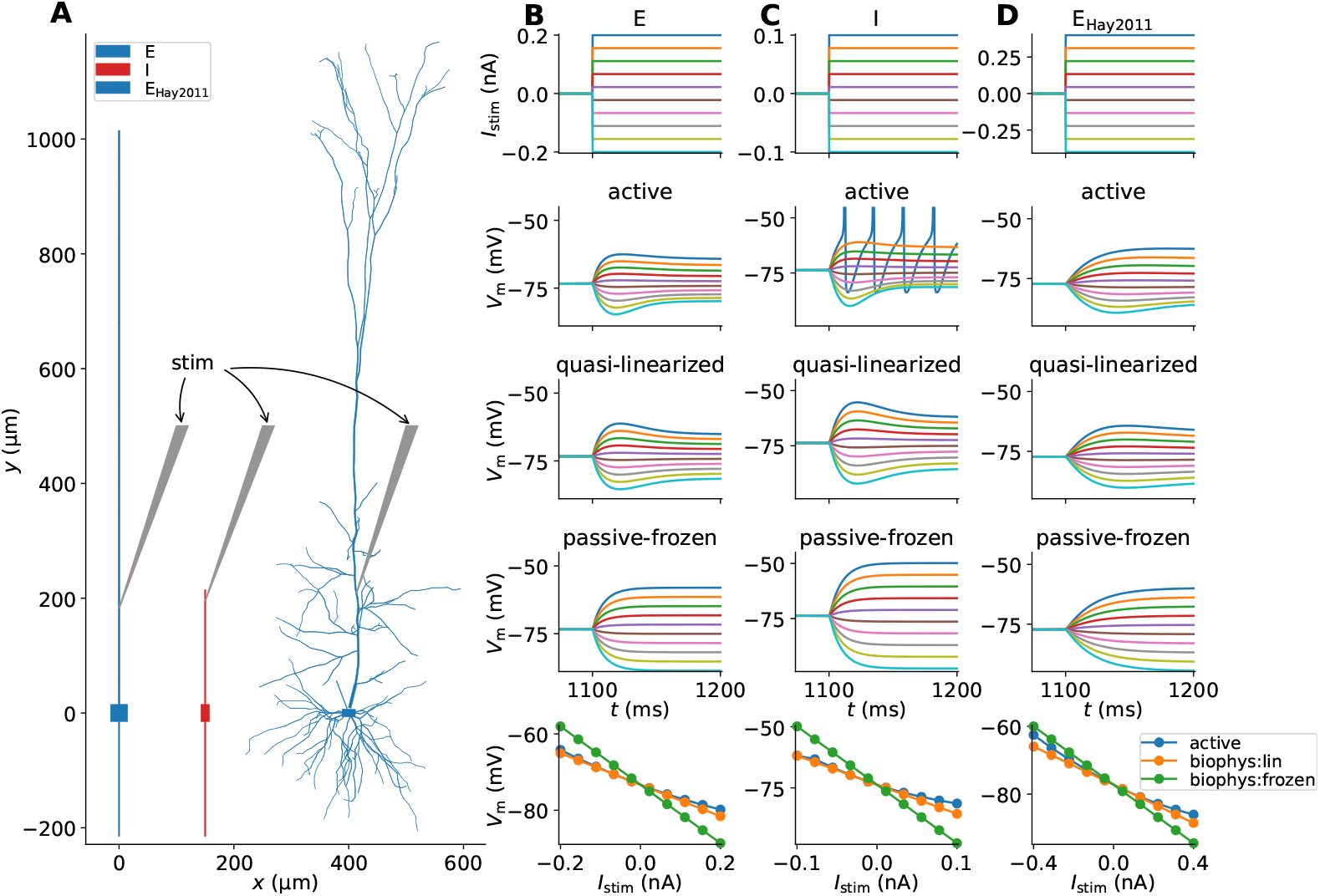
Active and linearized model neuron responses to dendritic current input. Same as Fig 2, but with current input in the apical dendrites approximately 200 μm from the soma compartments of the respective neurons.

**S2 Fig.**
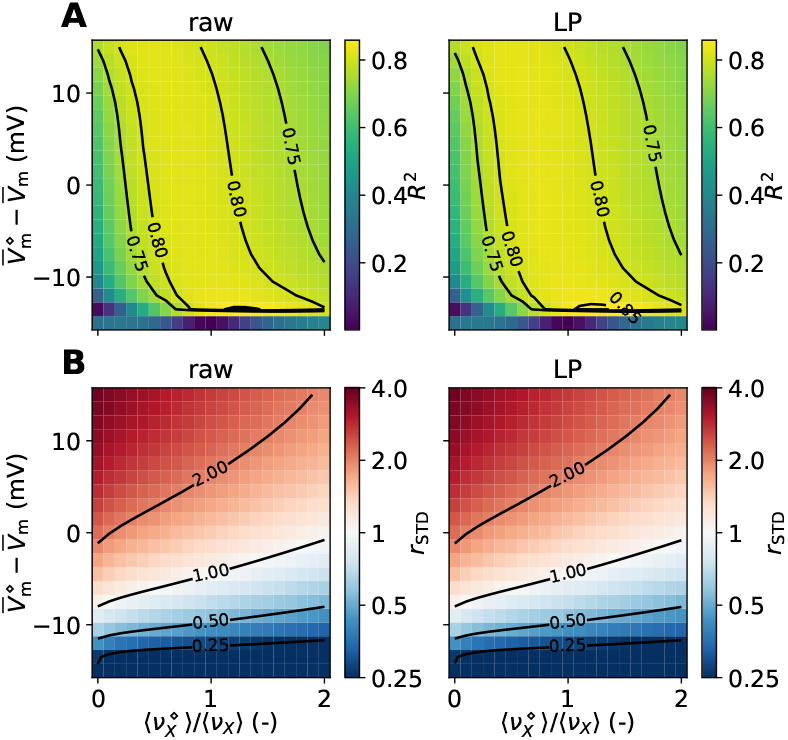
Effect of mismatched presynaptic firing rates and membrane potentials on kernel-based approximations to the current dipole moment signal. Same as Fig 8, but for the case where the excitatory (‘E’) population is replaced by biophysically detailed neuron models [30].

**S3 Fig.**
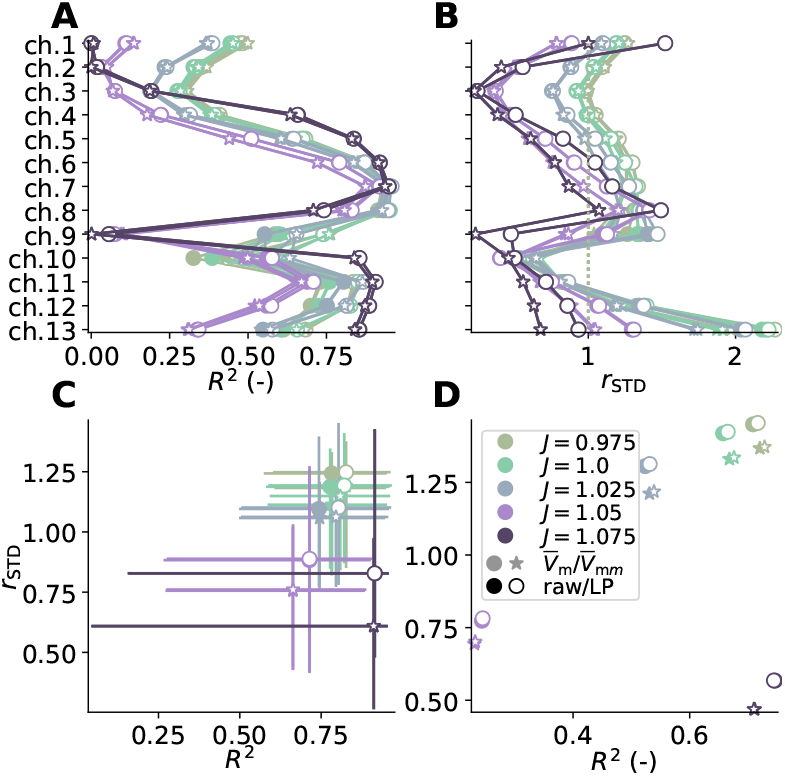
Effect of setting the linearization voltage 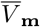 per segment. Same as Fig 13C-F, but for computed kernels and reconstructed signals either assuming a constant value for 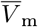 across the entire neuron model (circular lines/markers), versus kernel-based predictions where the 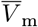 is set on a per-compartment basis (asterisk markers). For this, we use averaged values from each reference network simulation providing ground truth signals for comparison. Same color coding as in Fig 13.

## Acknowledgments

This work was supported by the European Union Horizon 2020 Research and Innovation Programme under Grant Agreement No. 785907 and No. 945539 Human Brain Project (HBP) SGA2 and SGA3 [EH,TVN,GH,PNB,CL,AM,GTE]. We also acknowledge the use of Fenix Infrastructure resources, which are partially funded from the European Union’s Horizon 2020 Research and Innovation Programme through the ICEI Project under the Grant Agreement No. 800858 [EH,SHM,TVN,GH,GTE]; The Helmholtz Alliance through the Initiative and Networking Fund of the Helmholtz Association and the Helmholtz Portfolio theme Supercomputing and Modeling for the Human Brain [PNB,CL,AM]; and The Excellence Strategy of the Federal Government and the Länder [G:(DE-82)EXS-PF-JARA-SDS005, G:(DE-82)EXS-SF-neuroIC002] [PNB,CL,AM]. The funders had no role in the study design, data collection, and analysis, decision to publish, or preparation of the manuscript.

## Authors contributions

EH, SMH, TVN, GH, and GTE conceived and conceptualized the project. EH drafted the manuscript. SMH, TVN, GH, PNB, CL, AM, and GTE co-wrote the manuscript. PNB and CL adapted and implemented the FIR filter implementation in NESTML. TVN implemented linearized active ion channels. EH wrote and ran all remaining simulation, analysis, and plotting codes for this paper.

## Footnotes

1 Contributions by presynaptic activity, that is, transmembrane currents of presynaptic neurons from APs and axonal propagation are not accounted for.

2 numba.pydata.org

3 cython.org

